# Harnessing molecular mechanism for precision medicine in dilated cardiomyopathy caused by a mutation in troponin T

**DOI:** 10.1101/2024.04.05.588306

**Authors:** Lina Greenberg, W. Tom Stump, Zongtao Lin, Andrea L. Bredemeyer, Thomas Blackwell, Xian Han, Akiva E. Greenberg, Benjamin A. Garcia, Kory J. Lavine, Michael J. Greenberg

## Abstract

Familial dilated cardiomyopathy (DCM) is frequently caused by autosomal dominant point mutations in genes involved in diverse cellular processes, including sarcomeric contraction. While patient studies have defined the genetic landscape of DCM, genetics are not currently used in patient care, and patients receive similar treatments regardless of the underlying mutation. It has been suggested that a precision medicine approach based on the molecular mechanism of the underlying mutation could improve outcomes; however, realizing this approach has been challenging due to difficulties linking genotype and phenotype and then leveraging this information to identify therapeutic approaches. Here, we used multiscale experimental and computational approaches to test whether knowledge of molecular mechanism could be harnessed to connect genotype, phenotype, and drug response for a DCM mutation in troponin T, deletion of K210. Previously, we showed that at the molecular scale, the mutation reduces thin filament activation. Here, we used computational modeling of this molecular defect to predict that the mutant will reduce cellular and tissue contractility, and we validated this prediction in human cardiomyocytes and engineered heart tissues. We then used our knowledge of molecular mechanism to computationally model the effects of a small molecule that can activate the thin filament. We demonstrate experimentally that the modeling correctly predicts that the small molecule can partially rescue systolic dysfunction at the expense of diastolic function. Taken together, our results demonstrate how molecular mechanism can be harnessed to connect genotype and phenotype and inspire strategies to optimize mechanism-based therapeutics for DCM.

**Significance statement:** Dilated cardiomyopathy (DCM), a leading cause of heart failure, is characterized by the inability of the heart to perfuse the body at normal filling pressures. There are multiple causes of DCM, including point mutations in sarcomeric proteins, but most patients receive similar courses of treatment, regardless of the underlying cause of the DCM. Many patients remain unserved by current therapies, and there is a need for new approaches. Here, we use multiscale experimental and computational approaches to demonstrate how knowledge of molecular mechanism can be harnessed to accurately predict the effects of a patient-specific mutation and responses to presumptive therapeutics. Our approach lays the foundation for a precision medicine approach to DCM.

## Introduction

Dilated cardiomyopathy (DCM), a leading cause of heart failure and indication for heart transplantation, is characterized by systolic dysfunction and dilation of the left ventricular chamber. DCM can be caused by a wide variety of pathologies, including ischemic heart disease, myocarditis, and genetic mutations in a wide variety of genes, including proteins associated with sarcomeric contraction (e.g., TTN, TNNT2, MYH7), gene expression (e.g., LMNA, RBM20), proteostasis (e.g., BAG3), and calcium handling (e.g., PLN) (1–3). Patients with DCM generally receive a similar course of treatment regardless of the underlying pathology driving the disease, and current guideline directed medical treatment does not universally improve outcomes in all patients (4). For example, for children with DCM, the 5-year transplant-free survival rate remains ∼50% using current therapies (5). Moreover, genotype positive DCM patients have reduced responses to current therapies compared to genotype negative patients (6). Thus, there is an outstanding need to develop tools and treatments that improve outcomes for patients with DCM.

Current treatments for DCM have focused on preventing adverse remodeling of the ventricles in broad patient populations without regard for the underlying pathology driving the disease pathogenesis (7, 8). An alternative approach would be to take a mechanism-based precision-medicine approach tailored to subgroups of patients. In this approach, subgroups of patients with common functional molecular defects would receive similar treatments tailored to reversing that defect (3). While a precision medicine approach holds potential promise for the future, it is not clear whether such an approach is feasible, in part because of challenges connecting genotype with phenotype. In particular, it remains challenging to understand the connections between point mutations and molecular and cellular dysfunction. This is especially difficult for patients in end-stage disease, since the molecular defect driving the disease pathogenesis often gets obscured by the complex adaptive and maladaptive remodeling accompanying heart failure.

To begin to bridge the chasm between genotype, the early disease pathogenesis, and potential mechanism-based treatments for DCM, we used multiscale experimental and computational tools to study a well-established mutation in troponin T, ΔK210 that causes DCM in patients (9–12). This mutation has been identified in at least 4 unrelated families as well as many sporadic cases, and several cellular and animal-based model systems of this mutation have been developed (13–19). These studies have clearly established the pathogenicity of this mutation for DCM. We previously investigated the molecular mechanism of this mutation, and we demonstrated that at the molecular scale, ΔK210 changes the positioning of tropomyosin along the thin filament, leading to reduced thin filament activation (13).

Here, we tested whether it is possible to use knowledge of molecular mechanism to computationally predict the early disease contractile phenotype in human cardiomyocytes and engineered heart tissues heterozygous for the mutation. We then leveraged this knowledge to identify a compound that partially reverses the molecular defect, and we show that it is possible to computationally predict both the beneficial systolic and less desirable diastolic responses of engineered heart tissues to the drug. Our results and approach lay out a practical pathway towards mechanism-based precision medicine for DCM.

## Results

### Harnessing molecular mechanism and computational modeling to predict compounds that will reverse reduced contractility with 11K210

The troponin complex, together with tropomyosin, regulates the calcium-dependent interactions between myosin and the thin filament (20). The process of thin filament activation depends on several factors, including calcium binding to troponin, movement of tropomyosin along the thin filament, and cooperative binding of myosin to the thin filament (**Fig. 1A**) (21). Previously, we showed that ΔK210 reduces thin filament activation by reducing the equilibrium constant between the closed and open states of the thin filament (13). We hypothesized that compounds that increase thin filament activation to reverse this molecular hypocontractility might be able to improve contractile function in human heart tissues.

**Figure 1:**
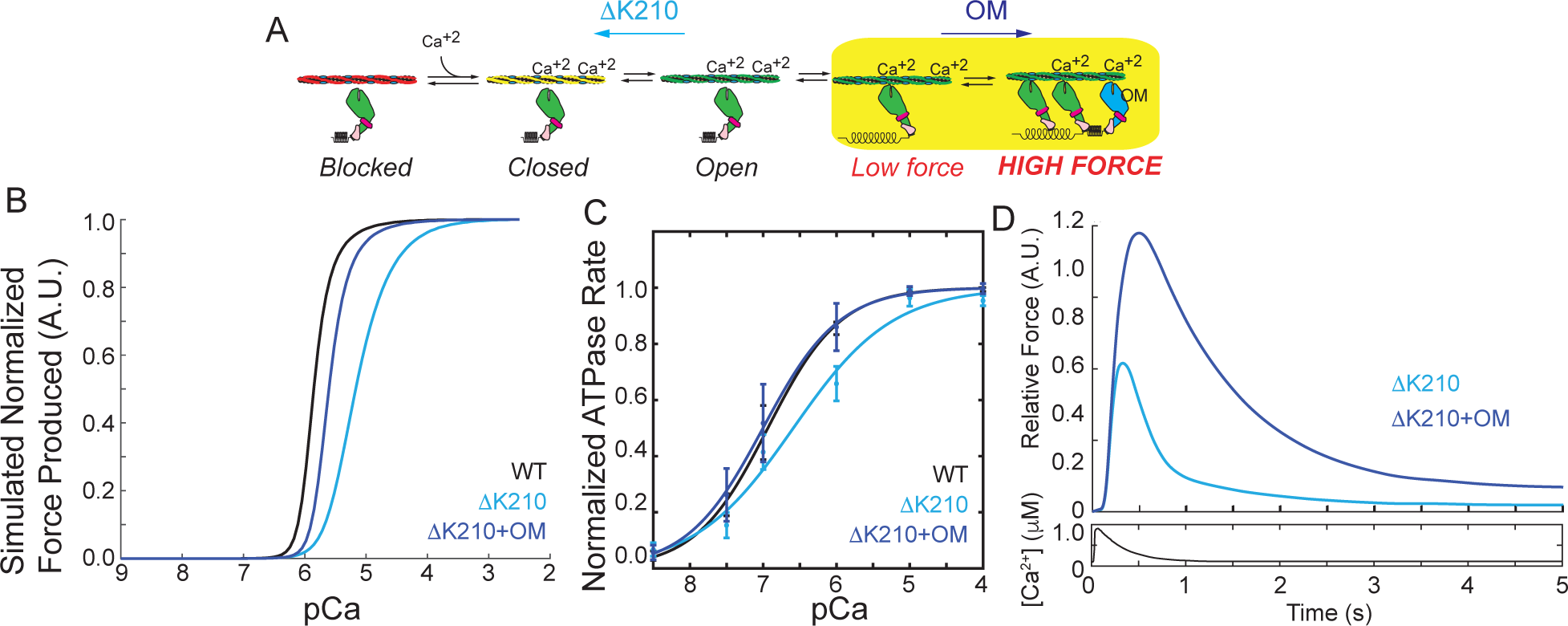
Computational modeling reveals potential therapeutic mechanisms. (**A**) Kinetic scheme showing the steps of thin filament activation. 11K210 reduces thin filament activation. Omecamtiv mecarbil (OM) increases cooperative activation of the thin filament by recruiting additional, non-force generating crossbridges. (**B**) Computational modeling of the normalized force produced as a function of calcium for WT and 11K210 based on the kinetic scheme in (A). 11K210 causes a shift towards supermaximal calcium activation, indicative of reduced thin filament activation. The model predicts that OM can partially rescue this shift by activating the thin filament. (**C**) Normalized ATPase measurements of myosin interacting with regulated thin filaments over a range of calcium concentrations. 11K210 causes a shift towards supermaximal calcium activation that can be partially rescued by OM treatment. (**D**) Computational modeling of the relative force per sarcomere in response to a calcium transient for 11K210 in the presence and absence of calcium. OM is predicted to cause an increase in force and prolongation of the force transient.

Here, we used a computational model of thin filament activation to predict the effects of the mutation on a heterozygous background. This mechanical model uses the rates and equilibrium constants underlying thin filament activation to predict both the steady-state and transient responses of the system to calcium (22). We assumed that the only difference between the WT and mutant proteins was the measured decrease in the equilibrium constants related to tropomyosin movement along the thin filament (13), and we modeled heterozygosity by assuming that half of the units are WT and half of the units are mutant. We then simulated the expected force per sarcomere over a range of calcium concentrations to generate a force-calcium curve. The simulation predicts that changing this parameter alone for the mutant is sufficient to shift the force per sarcomere curve towards supermaximal calcium activation (i.e., an increased pCa50, meaning that more calcium is necessary to reach half-maximal activation) (**Fig. 1B**). We experimentally tested this hypothesis by measuring the calcium-dependent ATPase rate of myosin interacting with thin filaments, and we see that ΔK210 causes a similar shift towards supermaximal calcium activation (**Fig. 1C**). This agrees well with our previous experimental measurements using an *in vitro* motility assay (13) and studies in other model systems (16, 18, 23, 24).

We hypothesized that compounds that reverse the defect of ΔK210 by increasing activation of the thin filament would rescue the molecular hypocontractility. Omecamtiv mecarbil (OM) is a myosin-binding compound that is currently in phase III clinical trials for systolic heart failure (25, 26). Biophysical studies have shown that OM has a complex biophysical mechanism (27–30) that includes 1) prolonging the amount of time that myosin remains bound to the thin filament while 2) inhibiting the force-generating myosin working stroke. It has been suggested that these long-lived crossbridges could keep the thin filament activated longer, leading to increased recruitment of other myosin heads.

We used our knowledge of OM’s biophysical mechanism (28) to computationally predict the effects of OM on thin filament activation for ΔK210. The model predicts that OM will shift the heterozygous mutant pCa50 closer to the WT, partially rescuing the molecular defect (**Fig. 1B**). We tested this experimentally using the ATPase assay, and we can clearly see that OM shifts the pCa50 of the mutant curve back towards the WT (**Fig. 1C**).

Next, we used the computational model to predict the response of the heterozygous mutant to a calcium transient. It has previously been established that OM does not appreciably affect the calcium transient (26), and therefore we used the same transient for simulating both treated and untreated conditions. The modeling predicts that OM treatment of the mutant will increase the force per sarcomere while simultaneously slowing relaxation (**Fig. 1D**). Taken together, our modeling predicts that ΔK210 reduces thin filament activation at the molecular scale, but OM might be able restore this activation. Moreover, the modeling predicts OM could improve systolic function in the mutant while prolonging activation, which may compromise diastolic function.

### 3D Engineered heart tissues hiPSC-CM^τιK210/WT^ show reduced systolic function

We then set out to test the predictions from our computational modeling in human engineered heart tissues (EHTs). The ΔK210 mutation was introduced into a human induced pluripotent stem cell (hiPSC) using CRISPR/Cas9, and we obtained several independent heterozygous cell lines (hiPSC^τιK210/WT^) as confirmed by next generation sequencing. The WT and heterozygous mutant cell lines are isogenic except for the 11K210 mutation, and whole exome sequencing showed that the parent line has no other known mutations associated with cardiomyopathies (13). To make EHTs, we generated both hiPSC derived cardiomyocytes (hiPSC-CMs) and hiPSC derived cardiac fibroblasts (hiPSC-CFs). For the hiPSC-CM differentiation, cardiomyocytes were enriched by metabolic selection (31). Using this procedure, we routinely obtain >90% cardiomyocytes as assessed by immunofluorescence staining for troponin T (TNNT2) (13, 32, 33). hiPSC-CMs were aged at least 30 days before conducting any experiments. We have validated the differentiation of hiPSC-CFs by qPCR measurement of key genes (34, 35).

Tissues were seeded in a matrix containing both collagen and Matrigel, as we have previously done (35). Incorporation of cells into EHTs provides additional cues found in the heart that are not present in 2D culture including multiple cell types, interactions with the extracellular matrix, and a three-dimensional geometry (1, 36). Moreover, incorporating hiPSC-CMs into EHTs promotes maturation of cells due to better electrical and mechanical coupling between cells. Here, we used two independent human engineered heart tissue (EHT) platforms to study the properties of hiPSC-CM^1′K210/WT^.

First, we generated linear tissue strips between two deformable polydimethylsiloxane (PDMS) pillars (**Fig. 2A**) (37). In this system, the tissues are grown under uniaxial tension to promote tissue organization. The active force of beating is calculated from the displacement and stiffness of the pillars. Tissues were aged for 14 days, and their contractile properties were measured under 1 Hz electrical stimulation. Both WT and mutant EHTs showed robust contraction under electrical stimulation (**Supplemental Movies 1 and 2**); however, the active forces generated by hiPSC-CM^1′K210/WT^ EHTs were significantly lower than the forces generated by the WT tissues (**Figs. 2B-D and Supplemental Figure 1**), consistent with our computational prediction (**Fig. 1D**).

**Figure 2:**
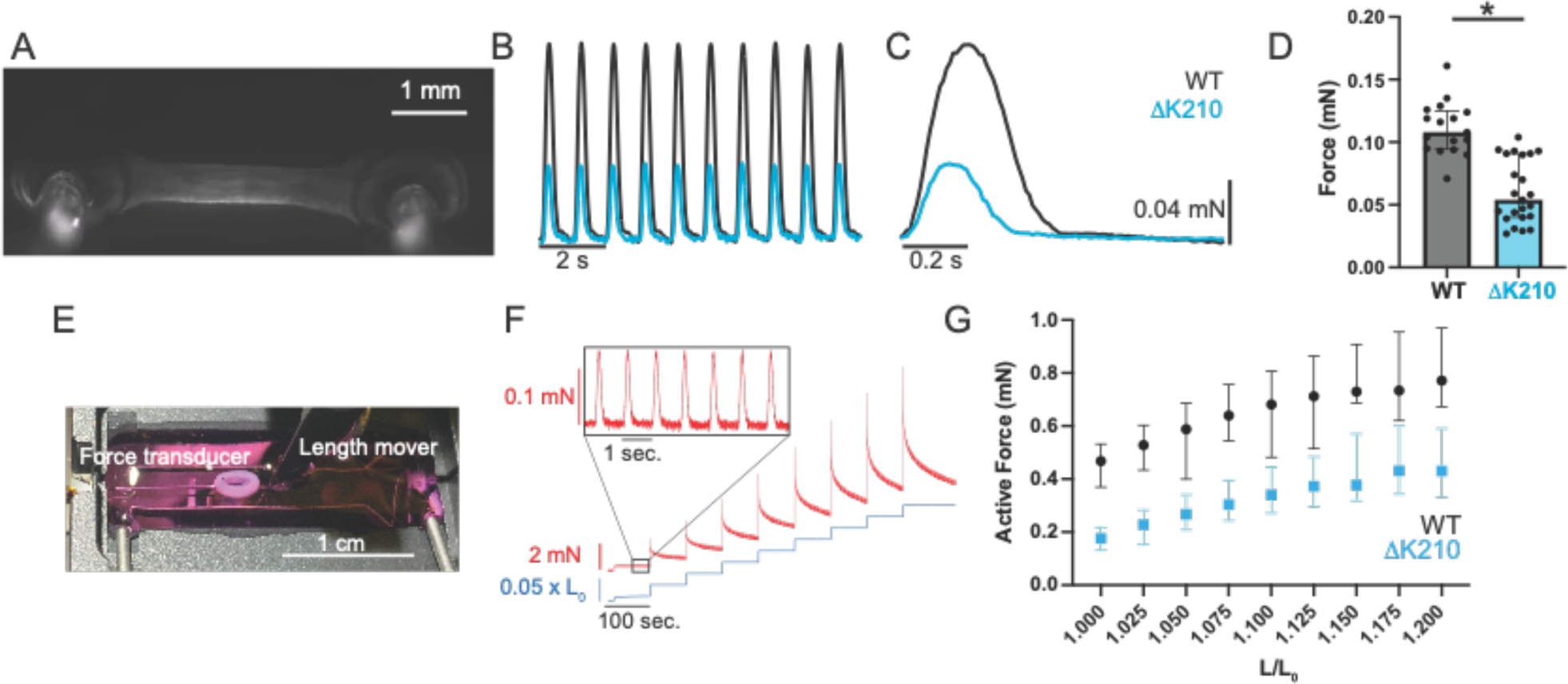
Engineered heart tissues with hiPSC-CM^1′K210/WT^ show reduced force production. (**A**) Linear engineered heart tissue grown between two deformable PDMS posts. (**B**) Representative data traces of WT and hiPSC-CM^1′K210/WT^ EHTs under 1 Hz stimulation. See Supplemental Movies 1 and 2. (**C**) Ensemble averaged beats for WT and hiPSC-CM^1′K210/WT^ EHTs shown above. (**D**) Data from N=17 WT and N=23 hiPSC-CM^1′K210/WT^ EHTs. Mutant 3D EHTs show reduced force production. Solid bars show median and error bars show interquartile range. See Supplemental Fig. 1 for additional parameters. (**E**) Ring-shaped EHT mounted between a length mover and a force transducer. (**F**) Representative trace showing the measured force of an EHT stimulated at 1 Hz undergoing a series of length steps. Inset shows the active force of beating. (**G**) Active force of beating as a function of stretch. Median values and 95% confidence intervals for N=11 WT and N=34 hiPSC-CM^1′K210/WT^ EHTs. The active force increases with stretch, consistent with the Frank-Starling relationship. A 2-way ANOVA shows significant differences between WT and hiPSC-CM^1′K210/WT^ EHTs with both stretch (P <0.001) and genotype (P<0.001), indicating that the hiPSC-CM^1′K210/WT^ EHTs show reduced force.

To further examine the length-dependent contractile properties of EHTs, we generated ring shaped tissues consisting of hiPSC-CMs and hiPSC-CFs (**Fig. 2E**) (38, 39). After maturation for 14 days, tissues were mounted between a length mover and a force transducer for mechanical testing. After mechanical preconditioning, tissues were subjected to a series of length changes (**Fig. 2F**). The active force increased with stretch, consistent with the well-established Frank-Starling relationship (**Fig. 2G**). While both WT and hiPSC-CM^1′K210/WT^ tissues showed increasing active force with stretch, the active force generated by hiPSC-CM^1′K210/WT^ tissues was significantly lower over the range of all stretches, consistent with our observation with the linear 3D tissue strips (**Fig. 2G**). Taken together, we see reduced contractility with hiPSC-CM^1′K210/WT^ tissues, consistent with our computational predictions and the observed reduction in contractility at the molecular scale (**Fig. 1**).

### hiPSC-CM^1′K210/WT^ show reduced force per sarcomere

While our experiments demonstrating reduced molecular contractility and our computational predictions of reduced force per sarcomere with the mutant are consistent with the reduced contractility seen in the EHTs, we wanted to probe additional mechanisms that could contribute to the observed reduction in force. Reductions in the force per sarcomere due to troponin dysfunction and/or reductions in the calcium transient or the number of sarcomeres in parallel in the cell due to downstream remodeling could all contribute to the reduced force seen in EHTs. We wanted to test the potential contributions of each of these non-exclusive possibilities at the single cell level, since it is challenging to probe these properties in the complex environment of a tissue.

We first examined the stress (i.e., force per area) generated by single hiPSC-CMs using traction force microscopy in which cells are patterned on to a deformable polyacrylamide substrate with embedded beads for tracking of contraction-induced substrate deformation. We generated substrates with 10 kPa stiffnesses that mimic the stiffness of the human heart, and cells were patterned on to rectangular islands of extracellular matrix to improve sarcomeric organization (13, 32, 33). Providing both stiffness and geometric cues has been shown to promote the maturation of cardiomyocytes (40). Both WT and hiPSC-CM^1′K210/WT^ cells responded to 1 Hz electrical stimulation and the traction stress was calculated for each cell (**Fig. 3A and Supplemental Fig. 2**). Consistent with our computational prediction and biochemical measurements, we observed that the hiPSC-CM^1′K210/WT^ cells generate less stress (i.e. force per area) than the WT cells.

**Figure 3:**
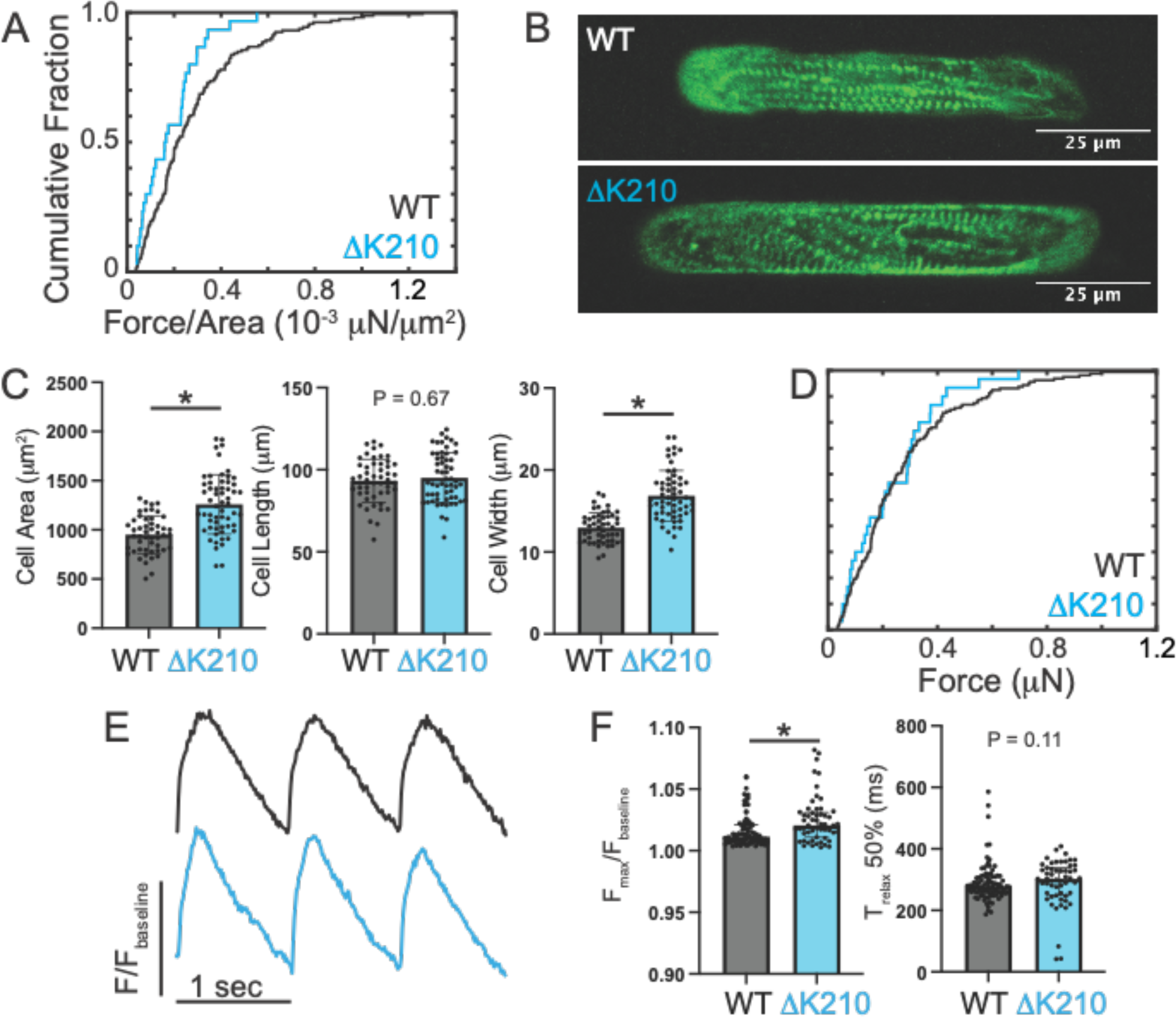
Measurements of single cell parameters. (A) Traction force microscopy of single hiPSC-CMs. Cumulative distribution showing the force per area for individual cells. N=159 WT and N=30 hiPSC-CM^1′K210/WT^ cells. The force per area is reduced in the mutant cells, as seen by a Kruskal–Wallis test (P<0.001). (**B**) Representative immunofluorescence confocal images of WT and hiPSC-CM^1′K210/WT^ cells patterned on to rectangular patterns of extracellular matrix. Sarcomeres are shown by alpha actinin antibody staining (green). (**C**) Quantification of cell size for N=50 WT and N=58 hiPSC-CM^1′K210/WT^. Solid bars show median and error bars show interquartile range. Statistical testing was done using a Mann-Whitney test. (**D**) Cumulative distribution showing the total force per cell in individual cells measured with traction force microscopy. A Kruskal–Wallis test was used to show that the force per area is not significantly different in the mutant cells (P=0.28). (**E**) Sample traces showing calcium transients for cells paced at 1 Hz. Signal was measured by fluorescence of FluoForte. (**F**) Quantification of calcium transients showing the amplitude of the transient above baseline (left) and the time to relax by 50% (right). N=91 WT and N=55 hiPSC-CM^1′K210/WT^ cells. Solid bars show median and error bars show interquartile range. Statistical testing was done using a Mann-Whitney test.

Next, we examined the size and morphology of hiPSC-CM^1′K210/WT^ cells using immunofluorescence (**Fig. 3B**). Microcontact printing was used to generate rectangular islands of extracellular matrix with a 7:1 aspect ratio for the hiPSC-CMs to improve maturation as described previously (13, 32, 33). Both WT and hiPSC-CM^1′K210/WT^ cells took on rectangular shapes when patterned; however, the hiPSC-CM^1′K210/WT^ cells were significantly larger than the WT cells (**Fig. 3C**). A similar increase in cell size was previously seen with the hiPSC-CM^1′K210/1′K210^ cells (13). We further examined the aspect ratio of these cells, and we saw that the hiPSC-CM^1′K210/WT^ cells show increases in width, but not length, consistent with concentric hypertrophy (**Fig. 3C**). The hypertrophy observed in 2D cultures with the mutant would be expected to increase the total force per cell. Therefore, we looked at the total force per cell from the traction force microscopy. Consistent with our prediction, we saw that while the force per area was lower in the mutant, the total force per cell was not significantly different in 2D culture (**Fig. 3D**). Interestingly, we did not observe increased cell size in the mutant EHTs (**Supplemental Fig. 3**), suggesting that adaptive increases in cell size highlighting key differences between 2D and tissue-like culture conditions.

Previously, it was shown that mutation and drug-induced reductions in force generation can alter sarcomeric organization (13, 41, 42). Therefore, we examined the sarcomeric organization by fixing the cells and staining for the sarcomeric Z-disc marker alpha actinin (**Fig. 3B**). The WT cells generally had well-formed sarcomeres that preferentially aligned along the long axis of the cells. While they also generally formed robust, aligned sarcomeres, hiPSC-CM^1′K210/WT^ sarcomeres showed more disorder, and this could contribute to the reduced force seen with the mutant. To test whether this disarray is related to the levels of expression of sarcomeric proteins, we measured the full proteomes of both WT and hiPSC-CM^1′K210/WT^ cells using mass spectrometry. Interestingly, hiPSC-CM^1′K210/WT^ EHTs show increased expression of key sarcomeric proteins including troponins I, T, and C (**Supplemental Fig. 4A**), and therefore, the reduced sarcomeric organization is not likely due to insufficient troponin levels. It is important to note that both WT and hiPSC-CM^1′K210/WT^ EHTs primarily express MYH7 myosin, and there is no difference in the levels between these groups (**Supplemental Fig. 4B**).

Finally, we examined the calcium handling of hiPSC-CM^1′K210/WT^ cells in 2D culture (**Fig. 3E**). Cells were loaded with the fluorescent calcium sensor FluoForte and stimulated at 1 Hz. The fluorescence was measured as a function of time using the CalTrack application (43). We found that the amplitude and kinetics of the calcium transient were only marginally affected in the hiPSC-CM^1′K210/WT^ cells (**Fig. 3F**). There is a slight (∼1%) increase in the amplitude of the calcium transient for hiPSC-CM^1′K210/WT^ cells, and this does not likely contribute to the reduction in contractility seen in EHTs.

Taken together, we demonstrate that at the cellular level hiPSC-CM^1′K210/WT^ cells show a reduction in the force per area, consistent with the experimentally measured reduction in molecular contractility and our computational predictions. Moreover, we do not observe downstream changes in cell size in the context of a tissue or changes in calcium handling that would contribute to the reduced force produced by hiPSC-CM^1′K210/WT^ EHTs. Finally, we observe changes in sarcomeric organization that could additionally contribute to the observed reduction in contractility.

### Reduced contractility in hiPSC-CM^1′K210/WT^ EHTs leads to transcriptional and proteomic remodeling of key proteins involved in mechanotransduction

At the molecular scale, the ΔK210 mutation in troponin T affects the functional properties of troponin, leading to reduced force production. Alterations in force can then lead to downstream changes in cellular processes via mechanobiological signaling pathways. Therefore, we investigated whether the mutation induces downstream transcriptional and proteomic remodeling in EHTs.

To look at transcriptional remodeling in the mutant, we performed RNAseq of both WT and mutant EHTs (**Fig. 4A and Supplemental Fig. 5**). We detected 22656 unique genes, with 3238 differentially expressed genes (log_2_fold change > 1, adjusted p-value < 0.05). KEGG pathway analysis on the differentially expressed genes revealed several pathways that are altered in the ΔK210 mutants including focal adhesion, extracellular matrix-receptor interactions, and cardiac muscle contraction **(Fig. 4B**). Significantly, several of these pathways have strong connections to mechanosensing and mechanotransduction. Therefore, we focused on the effects of the mutation on the extracellular matrix and focal adhesions. We found several key genes that have decreased expression in the mutant EHTs, including several integrin, laminin, and collagen isoforms (**Fig. 4C**). A heatmap of the top 50 differentially expressed genes in both genotypes can be found in **Supplemental Fig. 6**.

**Figure 4:**
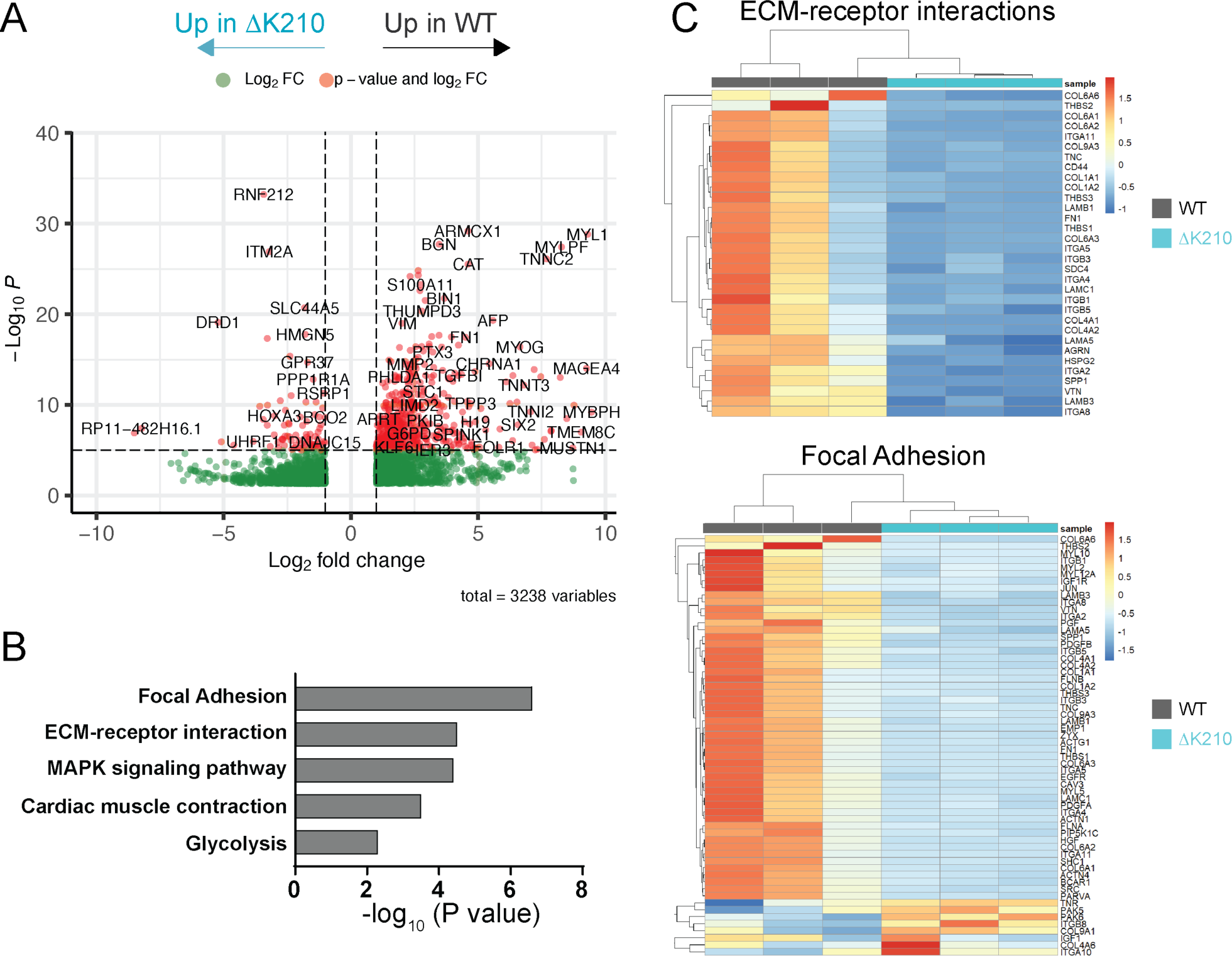
Whole transcriptome measurements for WT and hiPSC-CM^1′K210/WT^ EHTs. (**A**) Volcano plot showing differentially expressed genes between WT and hiPSC-CM^1′K210/WT^ EHTs. Cutoffs for differential expression are log_2_FC>1 and adjusted p-value < 10^-6^. (**B**) Kyoto Encyclopedia of Genes and Genomes (KEGG) terms associated with genes that are differentially expressed between WT and hiPSC-CM^1′K210/WT^ EHTs. Top terms include genes associated with extracellular matrix (ECM) receptor interactions and focal adhesions. (**C**) Heat maps showing differential expression of genes associated with ECM-receptor interactions and focal adhesions. Data is from N=3 different differentiations with multiple tissues pooled per sample.

To look at proteomic remodeling, we used mass spectrometry to view the whole proteomes of both WT and hiPSC-CM^1′K210/WT^ EHTs. We examined 7 independent tissues for each genotype from at least 2 separate differentiations. We detected 2595 unique proteins in our samples (**Fig. 5A**). Of these proteins, we saw 263 proteins that were differentially expressed (DE) between the WT and hiPSC-CM^1′K210/WT^ EHTs, with 144 proteins downregulated and 119 proteins upregulated in the mutant. When looking at the Gene Ontology (GO) cellular compartment terms associated with these proteins, we find that top terms include mechanotransduction pathways including cell-substrate junctions, focal adhesions, collagen-containing extracellular matrix, and the actin cytoskeleton (**Fig. 5B**). Differentially expressed proteins involved in mechanotransduction include laminin (LAMA4), dystrophin (DMD), filamin (FLNA, FLNB), and talin (TLN2) (**Fig. 5C**). Taken together, our results show that mutation-induced reductions in tension lead to downstream changes in the expression of key proteins associated with mechanotransduction.

**Figure 5:**
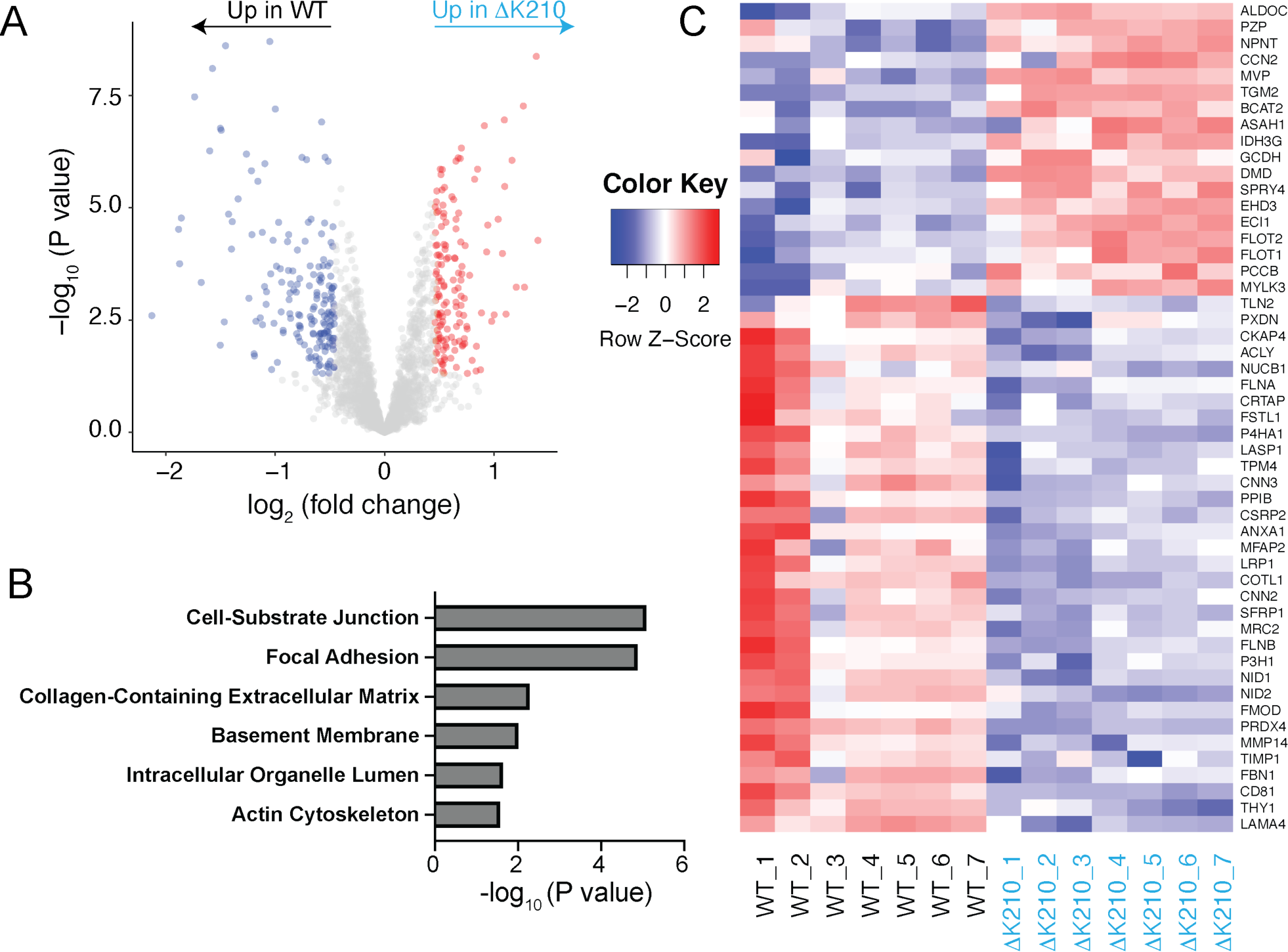
Whole proteome mass spectrometry measurements for WT and hiPSC-CM^1′K210/WT^ EHTs. (**A**) Volcano plot showing differentially expressed proteins between WT and hiPSC-CM^1′K210/WT^ EHTs. (**B**) Gene ontology (GO) terms associated with proteins that are differentially expressed between WT and hiPSC-CM^1′K210/WT^ EHTs. (**C**) Heat maps showing differential expression of key genes associated with ECM-receptor interactions and focal adhesions. Data is from N=7 different sample preparations with multiple tissues pooled per sample.

### Utilizing molecular mechanism to identify a compound that can rescue systolic function

Next, we set out to experimentally test whether knowledge of molecular mechanism can be harnessed to predict the effects of drugs on tissue function. As described above, our biochemical and computational studies predict that OM might be able to rescue the reduced thin filament activation seen with the mutant (**Fig. 1**). Therefore, we set out to test whether OM might be able to partially rescue the reduced systolic function seen with hiPSC-CM^1′K210/WT^ EHTs.

First, we tested whether acute treatment with OM (i.e., 20 minutes) could increase tissue contractility. The active force generated by linear EHTs was measured, then the EHTs were incubated with 250 nM OM for 20 minutes and the forces generated by the tissues measured again (**Fig. 6A**). 250 nM OM is similar to the therapeutic dose in patients (44–46). Consistent with our computational predictions (**Fig. 1D**), treatment of hiPSC-CM^1′K210/WT^ tissues with 250 nM OM significantly increased the force production (**Fig. 6A**). Next, we investigated the effects of chronic OM treatment on ring-shaped tissues. Tissues were treated with either 250 nM OM or DMSO for 5 days. Consistent with the computational modeling, treatment of mutant EHTs with OM increased the active force compared to DMSO controls (**Fig. 6B**).

**Figure 6:**
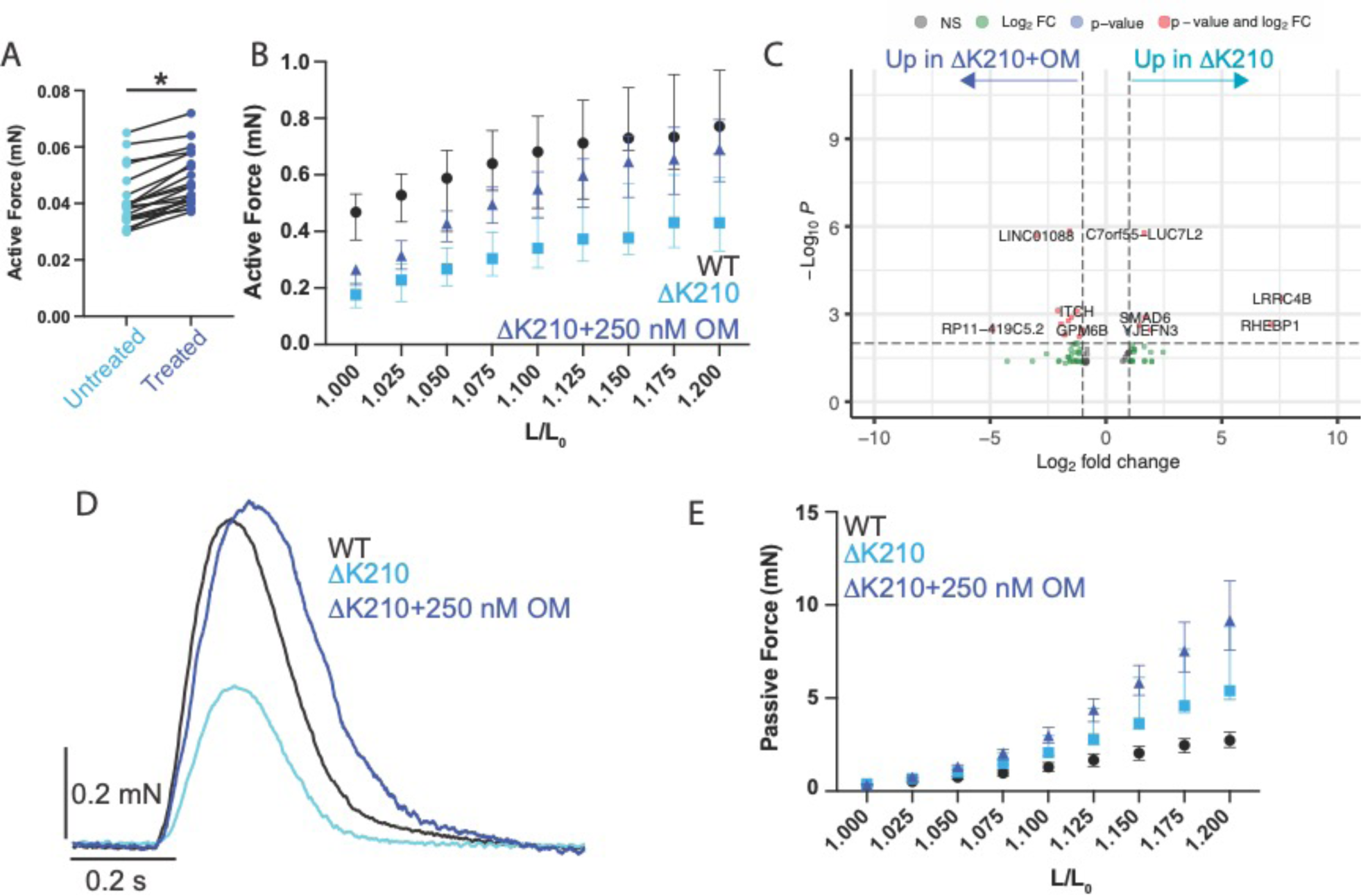
Mechanism-based rescue of contraction. (**A**) Linear hiPSC-CM^1′K210/WT^ EHTs acutely treated with 250 nM OM for 20 minutes. N=20 tissues. The force increased with treatment, as indicated by a Wilcoxon Matched Pairs Signed Rank Test (P<0.001). (**B**) Active force produced by tissues. Ring shaped tissues treated with 250 nM for 5 days (N=26 tissues). Shown are the median and 95% confidence intervals. A 2-way ANOVA shows significant differences in active force generation between WT, hiPSC-CM^1′K210/WT^ + DMSO, hiPSC-CM^1′K210/WT^ + 250 nM OM EHTs with both stretch (P<0.001) and genotype (P<0.001). hiPSC-CM^1′K210/WT^ + DMSO tissues generate less force than the WT over the range of all stretches (P<0.001). The force produced by the OM treated hiPSC-CM^1′K210/WT^ EHTs is not significantly different from the WT for stretches > 1.050 L/L_0_ (P>0.05). (**C**) Volcano plot showing differentially expressed genes between hiPSC-CM^1′K210/WT^ and hiPSC-CM^1′K210/WT^ + 250 nM OM EHTs. Cutoffs for differential expression are log_2_FC>1 and adjusted p-value < 10^-6^. 82 genes were identified. (**D**) Representative ensemble averaged trace showing the active force produced for hiPSC-CM^1′K210/WT^ tissues treated with DMSO or OM. Treatment with OM increased force and prolonged relaxation. For quantification, see Supplemental Figure 10. (**E**) Passive force produced by tissues after 5 days of treatment with 250 nM OM. Shown are the median and 95% confidence intervals. A 2-way ANOVA shows significant differences in passive force between WT, hiPSC-CM^1′K210/WT^ + DMSO, hiPSC-CM^1′K210/WT^ + 250 nM OM EHTs with both stretch (P<0.001) and genotype (P<0.001). All 3 conditions are significantly different from each other at stretches > 1.150 L/L_0_ (P<0.01, corrected for multiple comparisons).

While OM can partially rescue the observed contractile phenotype, we wanted to test whether partial reversal of the contractile phenotype is sufficient to reverse the downstream transcriptional remodeling seen in the mutant. Therefore, we performed RNAseq of WT and mutant EHTs treated for 5 days with 250 nM OM. We found that OM treatment had little effect on the transcriptional differences between the WT and mutant (**Fig. 6C and Supplemental Fig. 7-9**). In fact, many of the same differentially expressed mechanotransduction genes between the WT and the mutant remained affected, even after treatment with OM (**Supplemental Fig. 9**) This suggests that 5 days of treatment with OM can partially reverse the contractile phenotype, but that this is not sufficient to reverse the downstream transcriptional remodeling.

### OM treatment of hiPSC-CM^1′K210/WT^ EHTs affects diastolic function

In patients, OM causes impaired relaxation and diastolic function (47, 48). As described above, our computational modeling predicted that while OM would increase contractility in the mutant, it would also impair relaxation (**Fig. 1D**). To test this prediction, we looked at the ring-shaped EHTs and measured the rates of relaxation of hiPSC-CM^1′K210/WT^ EHTs treated for 5 days with DMSO or 250 nM OM. We clearly observe significantly prolonged time to relaxation in the OM treated tissues (**Fig. 6D and Supplemental Fig. 10**), consistent with our computational modeling.

Moreover, OM treatment increases the number of non-force generating crossbridges, which could increase tissue stiffness. To test whether OM affects the passive stiffness of the tissue, we further examined our measurements of the length-tension relationship for the ring-shaped EHTs treated with either DMSO or 250 nM OM for 5 days. Immediately after a length change, the tissues underwent viscoelastic relaxation to a steady-state force that represents the passive tension exerted by the tissue. The passive tension increases with increasing stretch, and the relationship between the stretch and the passive tension provides a measurement of the tissue stiffness. We found that treatment with OM increased the passive force on the hiPSC-CM^1′K210/WT^ tissues (**Fig. 6E**). Taken together, our results show that OM increases systolic function at the cost of diastolic function.

## Discussion

Here, we used computational modeling and multiscale experimental models to connect genotype and the early disease phenotype for a well-established DCM mutation, 11K210 in troponin T. We then leveraged our knowledge of molecular mechanism to predict the effects of a potential small molecule therapeutic on both systolic and diastolic function.

### Mutation-induced molecular dysfunction affects mechanics, driving the early disease pathogenesis

To better understand the initial connections between molecular and cellular dysfunction in the human disease, we developed hiPSC-CM^1′K210/WT^ cells to model the early disease pathogenesis. These cells are developmentally immature and thus they cannot recapitulate late-stage symptomatic disease and heart failure seen in patients and in animal models (16); however, they enable the modeling of the early disease pathogenesis in human cells (49). Subtle cardiac abnormalities can be observed in asymptomatic carriers of DCM mutations before the onset of pathological ventricular remodeling since the mutant protein is expressed throughout the lifetime of the patients (50, 51). As such, hiPSC-CMs are ideal tools for studying the early disease pathogenesis and for linking molecular and cellular dysfunction before the onset of pathological remodeling.

To experimentally model the early disease pathogenesis, we incorporated hiPSC-CMs into human engineered heart tissues. Importantly, culturing these cells in 3D tissues provides them with several biophysical cues that promote maturation and better mimic the environment of the heart including electrical coupling between cells, growth under tension, cellular interactions between cardiomyocytes and cardiac fibroblasts, and the incorporation of extracellular matrix proteins (36, 52). Providing these cues has been shown to promote maturation of hiPSC-CMs. Finally, our system used all human cells, which is important due to multiple physiological differences between human and non-human cells including their electrophysiology, calcium handling, and contractile protein isoforms (53) that can affect the disease presentation in different model systems (54).

At the molecular scale, point mutations affect the structure, function, and/or abundance of the protein (3). This is the initial molecular insult that drives the activation of adaptive and maladaptive pathways in DCM’s complex and protean disease pathogenesis. Previously, we showed that the fundamental molecular dysfunction caused by 11K210 is reduced thin filament activation, which reduces molecular contractility (13). Here, we used a well-established computational model of molecular contractility to predict the effects of the mutation on sarcomere-scale contractility on a heterozygous background. Our modeling predicted that incorporation of the mutant protein would cause a shift in the force-calcium curve towards supermaximal calcium activation, consistent with previous studies in other model systems (13, 16, 18, 23, 24) and our ATPase measurements (**Fig. 1C**). The modeling further predicts that this initial insult of reduced thin filament activation will lead to reduced force per sarcomere in response to a calcium transient, and we see this experimentally in both EHTs and single cardiomyocytes (**Figs. 2 and 3**). Thus, mutation-induced troponin dysfunction directly leads to altered contractility in cells and tissues.

While our data clearly reveals that ΔK210 directly affects molecular-based contractility, our transcriptomic and proteomic data clearly reveal that this molecular dysfunction drives downstream changes in cellular function and signaling (**Figs. 4 and 5**). This is consistent with previous studies showing that altered molecular mechanics can lead to downstream alterations in cellular and tissue function (24, 32, 33, 42, 55, 56). Several cellular structures have been identified to play roles in cardiomyocyte mechanobiology (57) including the matrix and integrins (58–60), the microtubule network (61), the nucleus (62, 63), and mecanosensitive signaling proteins such as YAP/TAZ (64, 65). Our transcriptomic and proteomic analyses revealed significant changes in extracellular matrix proteins and integrins that play critical roles in mechanotransduction in the heart (58–60). We speculate that mutation-induced changes in sarcomeric tension affect the tension transmitted between the extracellular matrix and nucleus, leading to altered sarcomeric organization and signaling. Consistent with this notion, cardiomyopathy mutations that affect force production, drug-induced changes in contractility, and genetic manipulation of proteins involved in linking the cytoskeleton to the extracellular matrix can all cause altered sarcomeric organization (13, 33, 41, 42).

### Mechanism-based correction of the underlying molecular defect

Since the ΔK210 mutation reduces thin filament activation (**Fig. 1**), we tested whether compounds that increase thin filament activation would improve systolic function. We examined omecamtiv mecarbil (OM), which was originally thought to be a myosin activator; however, subsequent biophysical studies have shown that the compound has a more complex mechanism of action (26–28), where OM-bound myosin crossbridges do not undergo a force-generating working stroke, rather, they remain strongly bound to the thin filament for a prolonged time (28). This increases the cooperative recruitment of other crossbridges, increasing thin filament activation and force production. Our computational modeling of thin filament activation is consistent with this notion (**Fig. 1**), and we can clearly see a shift in the pCa50 with OM in our ATPase measurements (**Fig. 1C**). At the same time, OM increases the population of non-force generating crossbridges, and our modeling predicts that this would prolong the time that the thin filament is activated (**Fig. 1D**). Taken together, our modeling predicts that OM would increase systolic function while decreasing diastolic function.

Here, we tested these model predictions in human engineered heart tissues. We found that both acute (**Fig. 6A**) and chronic treatment (**Fig. 6B**) of hiPSC-CM^1′K210/WT^ EHTs with OM increased force production as hypothesized. However, at the same time, we found that OM increased the stiffness of the tissues and prolonged systole (**Figs. 6D-E and Supplemental Fig. 10**), consistent with impaired relaxation (45, 47). This worsening of diastolic function was not due to changes in transcription or activation of fibroblasts, as we saw minimal changes in gene expression with OM treatment (**Fig. 6C**). Rather, both the improvement in systolic function and worsening of diastolic function could be predicted from knowledge of molecular mechanism.

While our study demonstrated that OM can partially rescue the systolic dysfunction caused by a thin filament mutation, it is important to note that increasing thin filament activation might not be ideal for all DCM mutations. The second most frequently mutated gene in DCM is the intermediate filament protein Lamin A/C (LMNA) (2). This protein localizes to the inner nuclear membrane where it plays roles in the regulation of gene expression and mechanotransduction; however, it does not play a direct role in sarcomeric contraction. As such, LMNA mutations have a different initial molecular insult, and the observed changes in contractility with LMNA mutations are likely secondary to that insult (66–69). Recently, it was shown that it is possible to rescue aspects of the LMNA DCM phenotype by targeting platelet derived growth factor receptor; however, this approach wouldn’t necessarily work with sarcomeric mutations (67). Similarly, we speculate that while treatment of LMNA mutants with OM could improve the contractile dysfunction, the disease phenotype might not be rescued since the LMNA-induced disruptions to mechanobiology and gene expression would remain. Our data imply that knowledge of molecular mechanism could be considered to identify populations most likely to respond to a given therapeutic intervention in clinical trials.

This study also highlights the need for new compounds for treating cardiomyopathy. Several small molecules targeting sarcomeric proteins are currently in development; however, it is important to consider their effects on both systole and relaxation. The two compounds that have advanced the farthest in clinical trials are myosin-targeting molecules: OM and mavacamten (70). OM increases systolic function while impairing relaxation, and mavacamten decreases systolic function while improving relaxation (26, 71). There is an outstanding need for compounds that can independently affect systolic and diastolic function; however, this continues to be a challenge for the field.

### Limitations

This study has several limitations. Importantly, even with state-of-the-art tools for maturing stem cell derived cardiomyocytes, these cells remain immature compared to adult cardiomyocytes (72). As such, they provide insights into the early disease pathogenesis, but extrapolation to treatment of symptomatic disease after the onset of pathogenic remodeling is a challenge. Our study, like all model system studies, gives insights into a specific time point (i.e., early disease pathogenesis), but the disease is protean and progressive in nature (3). Moreover, in vitro cellular systems can model aspects of the disease pathogenesis; however, they lack the cues and compensatory systems that are seen in whole organism studies. It is worth noting that treatments working early in the disease pathogenesis prior to the onset of maladaptive remodeling may not lead to complete reverse remodeling and restoration of function in the later stages of the disease. Finally, while our computational modeling was based on thin filament activation and capable of predicting effects on thin filament activation, other factors not captured in the model are likely at play in the heart.

### Concluding remarks and future prospects

Here, we demonstrate how multiscale models and knowledge of molecular mechanism can be applied to the study of a DCM mutation. It is possible that this platform could be used to study additional mutations and to identify optimal precision medicine therapeutics.

## Acknowledgements

The authors acknowledge financial support from Washington University in St. Louis and the Institute of Materials Science and Engineering for the use of instruments and for staff assistance. The authors would also like to acknowledge the financial support provided by the National Institutes of Health (R01 HL141086 to M.J.G.; NS 111997, P01 CA196539, and R01 HD106051to B.A.G.; R01 HL138466, R01 HL139714, R01 HL151078, R01 HL161185, and R35 HL161185 to K.J.L.), National Science Foundation (CHE2127882 to B.A.G.), Washington University BMB Seed Grant (PJ000027587 to Z.L.), the Children’s Discovery Institute of Washington University and St. Louis Children’s Hospital (PM-LI-2019-829 M.J.G. and K.J.L. and CH-II-2015-462, CH-II-2017-628 to K.J.L.), and the Washington University Center for Cellular Imaging (WUCCI). K.J.L. is also supported by the Washington University in St. Louis Rheumatic Diseases Research Resource-Based Center grant (NIH P30AR073752), Leducq Foundation Network (#20CVD02), Burroughs Welcome Fund (1014782), Foundation of Barnes-Jewish Hospital (8038–88), and generous gifts from Washington University School of Medicine.

## Author contributions

Conception and oversight by M.J.G., B.A.G., and K.J.L.. Cellular and tissue experiments were performed by L.G., and W.T.S. Biochemical experiments were performed by L.G. and A.E.G.. RNAseq experiments were performed by A.L.B.. Mass spectrometry experiments were conducted by Z.L. and X.H.. The computational model was programmed by T.B. and modelling was performed by M.J.G.. All authors contributed to the analysis of the data. The first draft was written by M.J.G.. All authors contributed to the writing and/or editing of the manuscript.

## Materials and Methods

### Computational modeling

For modeling of thin filament activation, we used a well-established computational model (22) based on the McKillop and Geeves model of thin filament activation (21). Details can be found in the Supporting Materials.

### Protein expression and purification

Human cardiac troponin I, T, and C were recombinantly expressed in E. coli, purified, and complexed as previously described (13, 73–75). The steady-state actin-activated myosin ATPase was measured using an enzyme coupled assay as previously described (76). Details can be found in the Supporting Materials.

### 11K210 TNNT2 stem cells

Stem cells were derived from the BJ fibroblast line (CRL-2522, ATCC) by the Washington University Genome Engineering and Stem Cell Core. Differentiation to hPSC-CMs was done in monolayer culture by temporal modulation of WNT signaling as previously described (13, 77). Cells were aged at least 30 days before performing any assays. All assays were conducted with 2 independent clones and cells were derived from at least 2 independent differentiations. Linear human engineered heart tissue strips were generated as previously described (35). Ring shaped EHTs were prepared as described in (78). Ring-shaped engineered heart tissues were mounted in an Aurora Scientific Small Intact Muscle Apparatus between a length mover and a force transducer. Details can be found in the Supporting Materials.

### Statistics

Figures show the median and interquartile ranges. Data was tested for normalcy using the Shapiro-Wilk test. Normally distributed data was analyzed using an ANOVA followed by post-hoc two-tailed T-tests with correction for multiple comparisons using the Tukey multiple comparison test. For data that did not follow a normal distribution, the data was analyzed using the nonparametric Mann-Whitney test. For paired measurements, we used a Wilcoxon Matched Pairs Signed Rank Test. For the analysis of the stretching data, we used a 2-way ANOVA interaction model. * denotes P < 0.05.

