## Supporting Materials for "Harnessing molecular mechanism for precision medicine in dilated cardiomyopathy caused by a mutation in troponin T"

### **Detailed Supporting Methods**

*Modeling*

For modeling of thin filament activation, we used a well-established computational model (1) based on the McKillop and Geeves model of thin filament activation (2). We have previously used this model to predict the effects of mutations on a homozygous background (3, 4). Briefly, we simulated a ring of 10 sarcomeric units where the probability of switching between states is given by previously established rate and equilibrium constants and a coupling constant (1). Using this model, it is possible to calculate the relative force per sarcomere at both steady state calcium (to generate the force-pCa curve) as well as the response to a calcium transient. To model heterozygosity, we followed the modification of (5) to average over the different possible permutations of where WT and mutant sarcomeres were located.

For our modeling of the WT, we kept all parameters identical to the default, validated parameters (1). To model sarcomeric units with the ΔK210 mutation, we only adjusted the equilibrium constant between the inactive and open states to match the relative change in this constant that we previously measured in the mutant (3). To model OM, we only made 2 changes to the model: 1) to mimic prolonged attachment duration, we adjusted the rate of exit from the strongly-bound state to equal the relative change in this constant between DMSO and OM that we previously measured and (2) to mimic inhibition of the working stroke, we reduced the force generated by this subunit to 0 for OM-bound crossbridges (6). Since the dosage used in our EHT experiments is close to the $K_d$ (6), we simulated ½ of the subunits bound to OM.

*Protein expression and purification*

Human cardiac troponin I, T, and C were recombinantly expressed in E. coli, purified, and complexed as previously described (3, 4, 7, 8). Human cardiac tropomyosin was expressed recombinantly in E. coli and purified as previously described (3, 4, 7, 8). Porcine cardiac actin and myosin were purified from cryoground porcine ventricles as previously described (3, 4, 8, 9). Myosin sub-fragment 1 (S1) was prepared by proteolytic cleavage as previously described (3, 4, 8, 9).

*Actin-activated steady-state ATPase measurements*

The steady-state actin-activated myosin ATPase was measured using an enzyme coupled assay as previously described (10). Briefly, globular actin was dialyzed overnight into ATPase assay buffer containing 20 mM imidazole pH 7.5, 10 mM KCl, 2 mM $MgCl_2$, and 1 mM DTT to polymerize the actin, and the actin was stabilized by adding a 1.1x molar excess of phalloidin. Tropomyosin was reduced before experiments by heating at 56$^o$C for 5 minutes in the presence of DTT and aggregates were removed by ultracentrifugation as previously described. 1 $\mu$M S1 myosin, 3.5 $\mu$M phalloidin stabilized actin, 2 $\mu$M troponin, 1 $\mu$M tropomyosin, 2 mM Mg*ATP, 0.313 mg/mL NADH (Sigma N7410-15VL), 0.5 mM Phospho(enol) Pyruvate (Sigma P0564), 1000 U/mL Pyruvate Kinase (Sigma P9136), 200 U/mL Lactate Dehydrogenase (Sigma L1254) and the desired concentration of free calcium were mixed in ATPase assay buffer containing 2 mM EGTA and 4 mM NTA. The concentration of free calcium was calculated using MaxChelator (11). Equivalent concentrations of DMSO were included in control and OM

(SelleckChem) treated experiments. Experiments were conducted in 96 well plates at 30$^{o}$C with shaking in a BioTek Synergy H1M plate reader. The absorbance at 340 nm was measured for 10 minutes and the absorbance decreased linearly. The rate of change in the absorbance was measured over a range of calcium concentrations. Five technical repeats from at least 2 separate protein preparations were used. Data was collected with paired treated and untreated conditions to minimize variability. Data were fitted with a Hill equation, where the ATPase rate V is given by:

$$V = V_o - \frac{V_{max} - V_o}{1 + e^{-h*(x - pCa_{50})}} \qquad \text{Equation 1}$$

Where x is the concentration of free calcium, $V_o$ is the basal ATPase rate under fully inhibited conditions, $V_{max}$ is the rate at fully activating conditions, h is the Hill coefficient, and the $pCa_{50}$ is the calcium concentration at which half-maximal activation is achieved.

*Generation of ΔK210 TNNT2 stem cells*

Stem cells were derived from the BJ fibroblast line (CRL-2522, ATCC) by the Washington University Genome Engineering and Stem Cell Core using the CytoTune127 iPS 2.0 Sendai reprogramming kit (A16517, ThermoFisher). Whole exome sequencing of this stem cell line revealed no mutations associated with cardiomyopathies (3). Two independent stem cell lines heterozygous for the ΔK210 TNNT2 mutation were derived using CRISPR/Cas9 and the genotype was verified by next generation sequencing as previously described (3).

*Differentiation of stem cells to human pluripotent stem cell derived cardiomyocytes*

Stem cells were maintained in feeder free conditions on Matrigel (Corning) coated plates in StemFlex media (ThermoFisher) as previously described. Cells were passaged using 0.02% EDTA at least 4 times before differentiating. Differentiation to hPSC-CMs was done in monolayer culture by temporal modulation of WNT signaling as previously described (3, 12). The population of hPSC-CMs was enriched by metabolic selection as previously described (3, 13). Using this protocol, we have routinely obtained >90% hPSC-CMs as assessed by immunofluorescence staining for troponin T (3). Cells were aged at least 30 days before performing any assays. All assays were conducted with 2 independent clones, and cells were derived from at least 2 independent differentiations.

*Differentiation of stem cells to human pluripotent stem cell derived cardiac fibroblasts*

Stem cell derived human cardiac fibroblasts were derived from the same cell line used to derive the cardiomyocytes according to the method of Zhang et al. (14), as we have previously done (15). Briefly, stem cells were differentiated towards a mesoderm lineage by modulation of WNT signaling and then directed towards a fibroblast lineage by modulation of FGF, WNT, and TGFβ signaling. Cardiac fibroblasts were validated by measuring the expression levels of both cardiac fibroblast specific genes, GATA4 and TCF21 as well as general fibroblast genes COL1A1 and DDR2 by RT-PCR.

*Immunohistochemistry of single cells on glass substrates and measurement of cell size*

Stem cell derived cardiomyocytes were patterned on glass using microcontact printing as previously described (3, 4, 9). Briefly, PDMS stamps for microcontact printing were generated using standard microfabrication techniques. Stamps contained an array of 17x118 µm (total area of 2000 µm$^2$) rectangles since this patterning promotes maturation (16). Stamps were incubated with Geltrex (Thermofisher) and then patterns were transferred to PLL-g-PEG coated glass as previously described (3). Cardiomyocytes were singularized in 0.25% trypsin, seeded on to the patterns at a density of 500 cells/µL in RPMI-20 (RPMI-1640 with 20% FBS) with 10 µM Y-27632, and allowed to attach to the surface for 45 minutes. Patterned cells were cultured in RPMI-20 with Y-27632 for 48 hours and were subsequently cultured in RPMI/B27 supplement with insulin, with media changes every 3 days.

Stem cell derived cardiomyocytes were fixed for 15 minutes in 4% formaldehyde in phosphate-buffered saline (PBS), permeabilized with 0.1% Triton X-100 (Sigma) for 20 minutes, and then blocked for 1 hour using a blocking solution containing 3% bovine serum albumin (Gold Bio), 5% donkey serum (Sigma), 0.1% Triton X-100, and 0.02% sodium azide in PBS as previously described (3). The primary antib (15)odies were then allowed to incubate with the cells overnight at 4°C. The next day, cells were thoroughly washed in PBS before being incubated with secondary antibody for 1 hour. Fixed cells were imaged on a Nikon A1Rsi confocal microscope equipped with an Andor Zyla 4.2 Megapixel sCMOS camera (Washington University Center for Cellular Imaging). Z-stacks of cells with 60x magnification were recorded in sequential scanning mode. Cell size was measured in ImageJ (17) using Z-stack projections as previously described (3).

| Antibody | Product Number | Working Dilution | Host species | Vendor |
|---|---|---|---|---|
| Alpha-actinin | A7811/EA-53 | 1:500 | mouse | Sigma |

*Linear human engineered heart tissue strips*

Linear human engineered heart tissue strips were generated as previously described (15). Briefly, tissues were seeded in casting molds treated with 1% pluronic-F127 to block the surface from adhering to the seeded tissues. To seed the tissues, 1 mg/mL final concentration rat collagen I and a final concentration of 0.77 mg/mL growth factor reduced Matrigel (Corning: 354230) were mixed with cardiomyocytes and fibroblasts. Each tissue contained $10^6$ hPSC-derived CMs and 5% fibroblasts. Silicon racks were placed above the mixture in the casting mold and the matrix was allowed to polymerize for 2 h at 37°C before being covered with RPMI-20 with 5 µM Y-27632. The following day, the tissues were transferred into media containing DMEM High glucose (4 mg/mL, Fisher Scientific: 10-013-CV), 10% FBS (Millipore Sigma: F0926), 1% Non-Essential Amino Acids (Life Technologies: 11140050), 1% GlutaMAX Supplement (ThermoFisher: 35050061), and 1% Pen-Strep (Gibco: 15140122). Media was changed every other day and mechanical experiments were performed on day 14 post seeding.

*Mechanical measurements of linear human engineered heart tissue strips*

On day 14 post seeding, tissues were moved into the measuring system (EHT Technologies). The measuring system was maintained at 37 °C in 40% oxygen and 5% carbon dioxide. Tissues were stimulated at 1 Hz using a Myopacer with 10 V 15 ms biphasic pulses (IonOptix). The displacement of the PDMS posts was measured by video microscopy by the automated system at 100 Hz. The measuring system provided measurements of several physiological parameters including peak force and the kinetics of contraction and relaxation.

*Generation of ring-shaped engineered heart tissues*

Ring shaped EHTs were prepared as described in (18). Briefly, PDMS molds were made by preparing PDMS at a 1:10 ratio and pouring into a 35 mm dish containing two 10 mm outer diameter x 4 mm inner diameter Teflon washers. The PDMS molds were baked overnight at 65°C and the washers were removed using ethanol leaving behind a 10 mm diameter round mold with a 4 mm center stalk.

Tissues were seeded into the PDMS molds. Working on ice, rat collagen I (Corning: 354236; 1 mg/mL final concentration) was combined with equal parts 2x DMEM supplemented with FBS, and the pH of the collagen/DMEM mixture was neutralized with sodium hydroxide. hPSC-CMs and fibroblasts were dissociated from 2D cultures using 0.25% Trypsin-EDTA, resuspended in media containing RPMI-20 and 10 μM Y-27632, and combined with the collagen mixture on ice. Each tissue contained $10^6$ hPSC-CMs and 5% fibroblasts. The mixture was allowed to polymerize for 1 hour in the PDMS molds and then covered with RPMI-20 and 5 μM Y-27632 media. The following day the media was changed to DMEM High glucose (4 mg/mL), 10% FBS, 1% non-essential amino acids, 1% GlutaMAX Supplement, and 1% Pen-Strep. Tissues were aged at least 10 days before collecting data. DMSO or omecamtiv mecarbil were added 5 days prior to collecting data.

*Mechanical measurements of ring-shaped engineered heart tissues*

Ring-shaped engineered heart tissues were mounted in an Aurora Scientific Small Intact Muscle Apparatus between a length mover and a force transducer. The system includes a Real-Time Data Acquisition and Analysis module that enables fast feedback

(100 Hz). Tissues were perfused with Tyrode's buffer containing 2 mM $CaCl_2$ at 37 °C and the temperature was maintained using a temperature controller. Tissues were electrically stimulated at 1 Hz with 10 V 15 ms biphasic pulses using a Myopacer from IonOptix. Tissues were stretched to 1.25x the resting length followed by 5 minutes of preconditioning, where the tissues were held under 2.5 mN of constant tension using a feedback loop. Tissues were then pretensed to 0.5 mN force, and the starting length was measured. Tissues were then stretched over 8 steps at 1-minute intervals, each time elongating the tissue by increments of 0.025x the starting length. The length and force were measured at 100 Hz.

Data traces were analyzed using custom software written in Labview and Matlab. Cross sectional areas were not used since all tissues had the same number of cells. The passive force was measured as the baseline reached after stretching from an exponential fit. The active force was measured from the average force generated during beating during the last 5-10 seconds before a length change. Irregular beats outside of pulsing were omitted from this analysis. Ensemble averaged force traces were generated by synchronization on the electrical stimulation and the rate of force relaxation were calculated from these traces.

*Immunohistochemistry of linear human engineered heart tissues*

Tissues were clarified overnight in RapiClear 1.52 solution (SunJin Lab Co.) and mounted using RapiClear 1.52 solution. Wells were created for whole mount tissues using double sided sticky iSpacer 0.25 mm (catalog IS206) placed on standard glass slides. The spacers could be doubled up for thicker tissues. Coverslips measuring 24x40 mm

(VWR) were placed on top of the iSpacer and the tissues were imaged on an inverted confocal microscope. Cells were stained for DAPI and Wheat Germ Agglutinin (WGA) Alexa Fluor 647 (ThermoFisher: W32466).

*Traction force microscopy*

Traction force microscopy was performed on 10 kPa polyacrylamide hydrogels as previously described (3) based on the method of Ribeiro et al (16). Briefly, stem cell derived cardiomyocytes were seeded on to hydrogels with rectangular shaped Geltrex patterns as described above. Hydrogels contained 0.2 μm diameter fluorescent beads (Bangs Beads: FSSY002) for tracking of movement. Single, cells were imaged under 1 Hz 15-20 V electrical stimulation (IonOptix) in an environmentally controlled chamber (Tokai Hit) on a Nikon spinning disk confocal microscope equipped with a Yokagawa CSU-X1 variable speed Nipkow spinning disk scan head and Andor Zyla 4.2 Megapixel sCMOS camera (Washington University Center for Cellular Imaging). Traction force videos were analyzed in Matlab as previously described (3, 19).

*Calcium transients*

Cardiomyocytes were plated on glass bottom 35 mm dishes (MatTek Corporation) and allowed to recover for a week before being used in experiments. Cells were loaded with 2.5 μM FluoForte (Enzo Life Sciences) in RPMI/B27 media and incubated for 20 min at room temperature protected from light. The cells were washed twice in Tyrode's solution containing 1.8 mM $CaCl_2$, 135 mM NaCl, 4 mM KCl, 1 mM $MgCl_2$, 5 mM glucose, and 10 mM HEPES pH 7, and incubated for an additional 15 minutes at 37°C to deesterify the AM dye. Cells were imaged in an Okolab environmental control chamber using a 40x water objective and a Hamamatsu C13440 camera at 100 frames per second for 20 seconds. Data were analyzed using CalTrack to calculate the $F/F_{baseline}$ signal and time parameters (20).

*RNA sequencing*

Total RNA was extracted from WT and ΔK210 linear engineered heart tissues (4 tissues per replicate) . RNA integrity was determined using a 4200 Tapestation. Library preparation was performed with 10 ng of total RNA with a Bioanalyzer RIN score greater than 8.0. ds-cDNA was prepared using the SMARTer Ultra Low RNA kit for Illumina Sequencing (Takara-Clontech) per manufacturer's protocol. cDNA was fragmented using a Covaris E220 sonicator using peak incident power 18, duty factor 20%, cycles per burst 50 for 120 seconds. cDNA was blunt ended, had an A base added to the 3' ends, and then had Illumina sequencing adapters ligated to the ends. Ligated fragments were then amplified for 12-15 cycles using primers incorporating unique dual index tags. Fragments were sequenced on an Illumina NovaSeq-6000 using paired end reads extending 150 bases.RNA-seq reads were then aligned and quantitated to the Ensembl release 101 primary assembly with an Illumina DRAGEN Bio-IT on-premise server running version 3.9.3-8 software. Results were analyzed using the DESeq2 package in R.

*Mass spectrometry*

*Sample preparation for whole proteomic analysis*

Engineered tissues from WT and ΔK210 (8-10 ring tissues each replicate) groups were added 400 μL of 8 M urea in PBS, lysed, and denatured by probe sonication (Fisherbrand™ Model 120 Sonic Dismembrator, setting: 1 sec on, 2 sec off, 20% energy, 60 cycles) to yield a homogeneous solution. Protein concentration was determined by DC assay (Bio-Rad). For whole proteomic analysis, 100 μg of proteome was diluted in 40 μL of 8 M urea in PBS, reduced by 5 mM of TCEP with 30 min incubation at 37 °C, alkylated by 15 mM of iodoacetamide (IAA) with 30 min incubation at room temperature in the dark. The solution was diluted to 2 M urea by 50 mM ammonium bicarbonate in H2O, digested by trypsin (sequence grade, Promega) at 1:50 trypsin/protein ratio (*w*/*w*) with overnight (~12 h) incubation at 37 °C. The resulted peptide solution was acidified by formic acid at a final concentration of 5%. 1/5 (~20 μg) of the peptide solution was desalted, resuspended in 0.1% formic acid at ~100 ng/μL. 200 ng of each sample was injected for LCMS analysis in data-independent mode.

*Sample desalting*

Peptides were resuspended in 0.1% formic acid and desalted by in-house packed stage-tips. Stage-tips were manufactured by sealing five disks of C18 material (cat. No.: 2315, Empore, 3M Company) at the bottom of a P200 tip. C18 disks were cut by sample corers (cat. No.: 18035-01, Fine Science Tools). Stage-tips were equilibrated with 50 μL of methanol, 50 μL of 80% acetonitrile in H2O containing 0.1% formic acid (FA), and 50 μL of water containing 0.1% FA by centrifugation (1,000 x g, ~1-2 min). The sample was loaded to the stage-tip, centrifuged to flow through, and washed with 75 μL of water containing 0.1% FA. The stage-tip was transferred to a new collection tube, eluted by 75 μL of 80% acetonitrile in H2O containing 0.1% FA. The sample was dried by SpeedVac (SAVANT SVC100H Refrigerated Condensation Trap) under vacuum for 30-60 min at room temperature, stored at -80 °C.

*Whole proteomic analysis*

An LC-MS/MS system consisted of a Waters M5 UHPLC coupled to a ZenoTOF 7600 (Sciex) was used for peptide analysis in data-independent mode. Peptide samples were maintained at 7 °C on sample tray in LC. Separation of peptides was carried out on an Waters nanoEase M/Z Symmetry C18 Analytical Column, (5 μm, 100 Å, 300 μm X 150 mm) at room temperature with a mobile phase consisting of a linear gradient of A (0.1%FA in $H_2O$) and B (acetonitrile containing 0.1% FA) under the following conditions: 0 → 1 → 46 → 47 → 50 → 50.5 → 55 min, 2 → 2 → 32 → 80 → 80 → 2 → 2% B. The flow rate was set at 5 μL/min. The ZenoTOF 7600 system was operated using the OptiFlow TurboV ion source with a vertical microflow probe (1-50 μL/min electrode). The Zeno SWATH DIA method consisted of 85 variable-width SWATH DIA windows that spanned the Q1 mass range 399.5-903.5 Da. MS/MS accumulation times of 13 ms were used with Zeno trapping over the MS/MS mass range 400-1500 Da. Zeno SWATH DIA data were processed using Spectronaut (v18.1) with library-free searches (directDIA). The analysis of differentially expressed (DE) proteins was carried out based on our previous work (21). Briefly, protein quantification data from Spectronaut was log transformed, calculated for p values, and calculated for the mean and derived log2(fold change) values for each protein. Statistical significance was determined based on FDR cutoff (0.05) and log2(fold change) cutoff (2 s.d.). R script for DE analysis is publicly available to download from GitHub at https://github.com/BeckyHan/Greenberg_Lab.

*Statistics*

Figures show the median and interquartile ranges. Data was tested for normalcy using the Shapiro-Wilk test. Normally distributed data was analyzed using an ANOVA followed by post-hoc two-tailed T-tests with correction for multiple comparisons using the Tukey multiple comparison test. For data that did not follow a normal distribution, the data was analyzed using a Mann-Whitney test. For paired measurements, we used a Wilcoxon Matched Pairs Signed Rank Test. For the analysis of the stretching data, we used a 2-way ANOVA interaction model. * denotes $P < 0.05$.

### Supplemental Figures

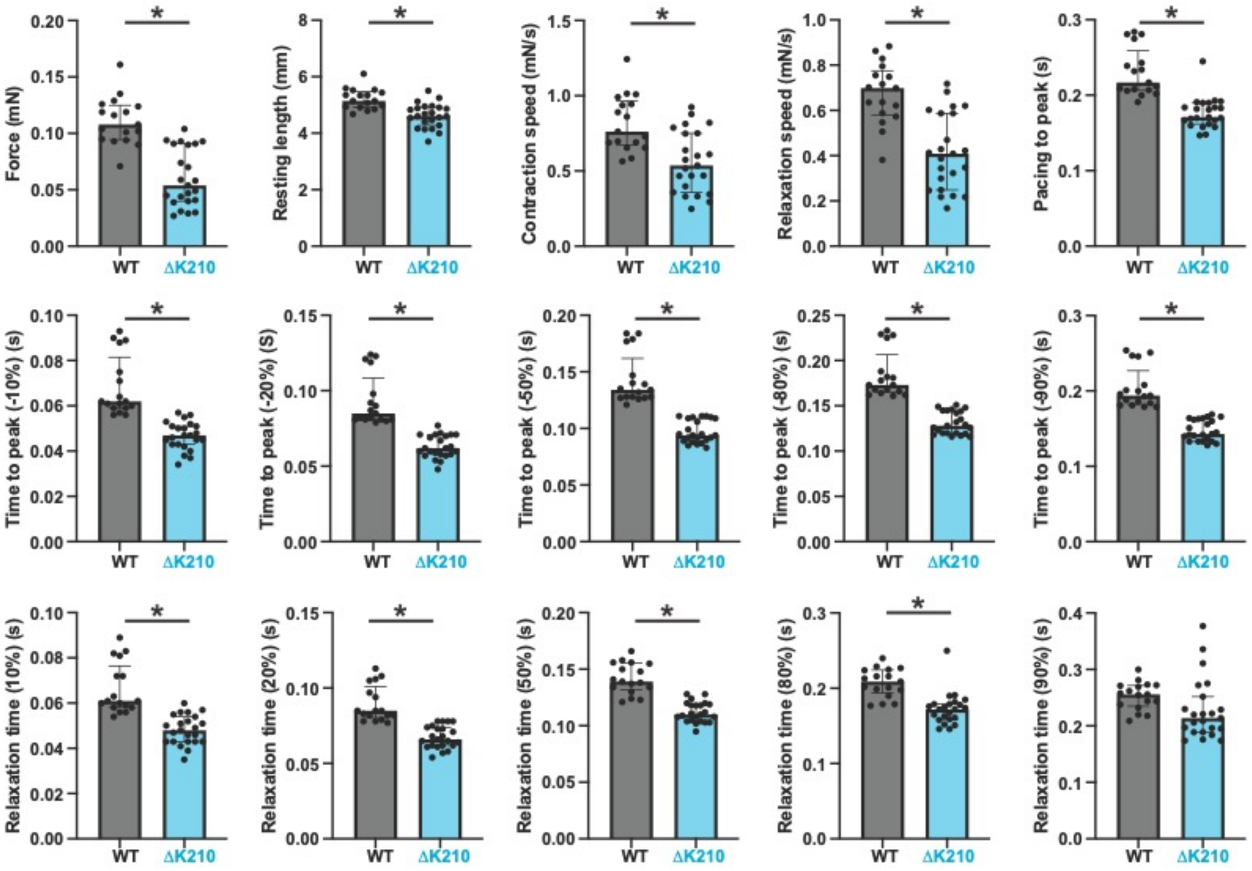


**Supplemental Figure 1: Key parameters measured for linear engineered heart tissues.** Data from N = 17 WT and N = 23 hiPSC-CM$^{\Delta K210/WT}$ EHTs. Mutant 3D EHTs show reduced force production. Solid bars show median and error bars show interquartile range. Statistical testing done by a Mann-Whitney test.

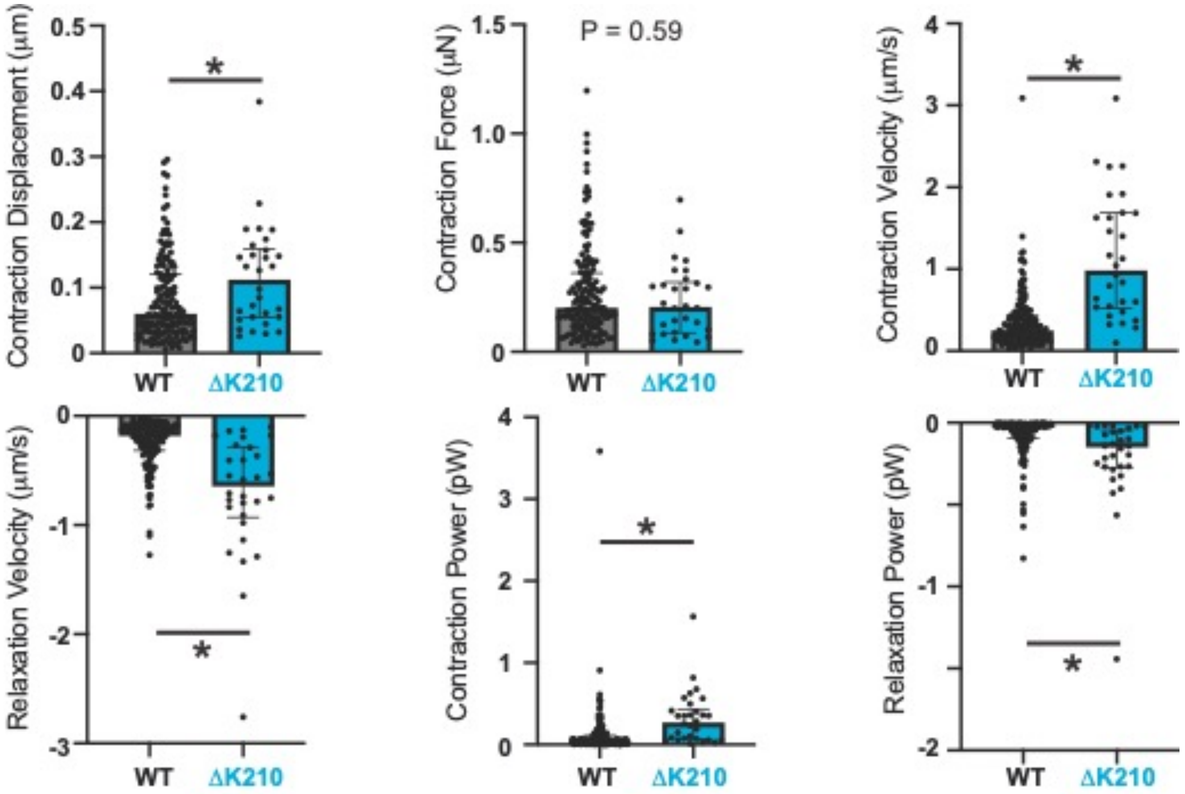


**Supplemental Figure 2: Key parameters measured by traction force microscopy for single cells.** Data from N = 159 WT and N = 30 hiPSC-CM$^{\Delta K210/WT}$ cells. Solid bars show median and error bars show interquartile range. Statistical testing done by a Mann-Whitney test.

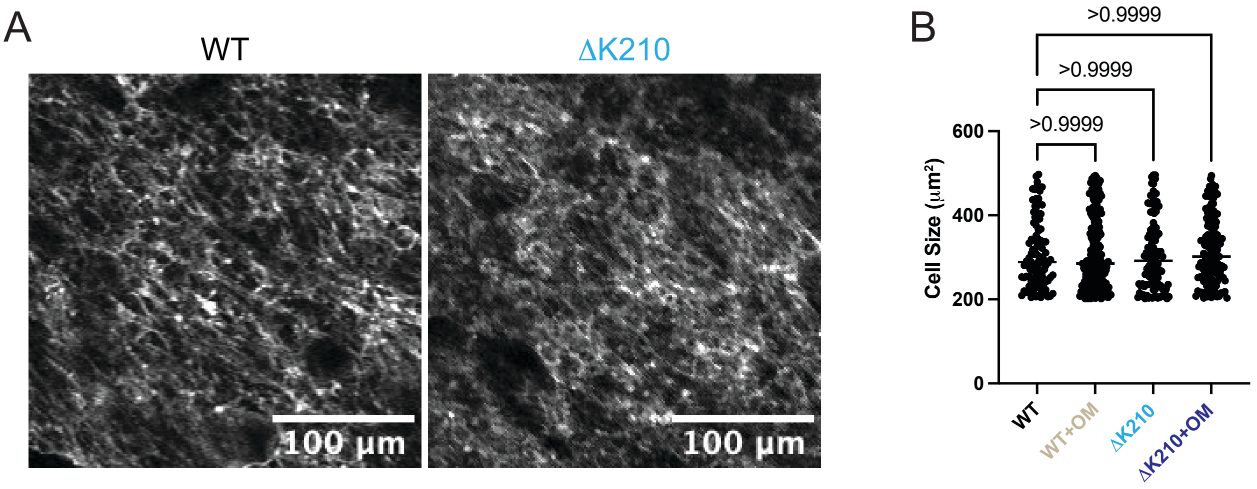


**Supplemental Figure 3: Fixed and stained linear EHTs for measurement of cell size.** (**A**) Representative images showing WGA stained EHTs. Cells were identified by thresholding in ImageJ (see methods). (**B**) Quantification of cell size. N = 104 WT + DMSO, N = 187 WT + OM, N = 114 ΔK210 + DMSO, and N = 133 ΔK210 + OM. A Mann-Whitney test was used for statistical testing.

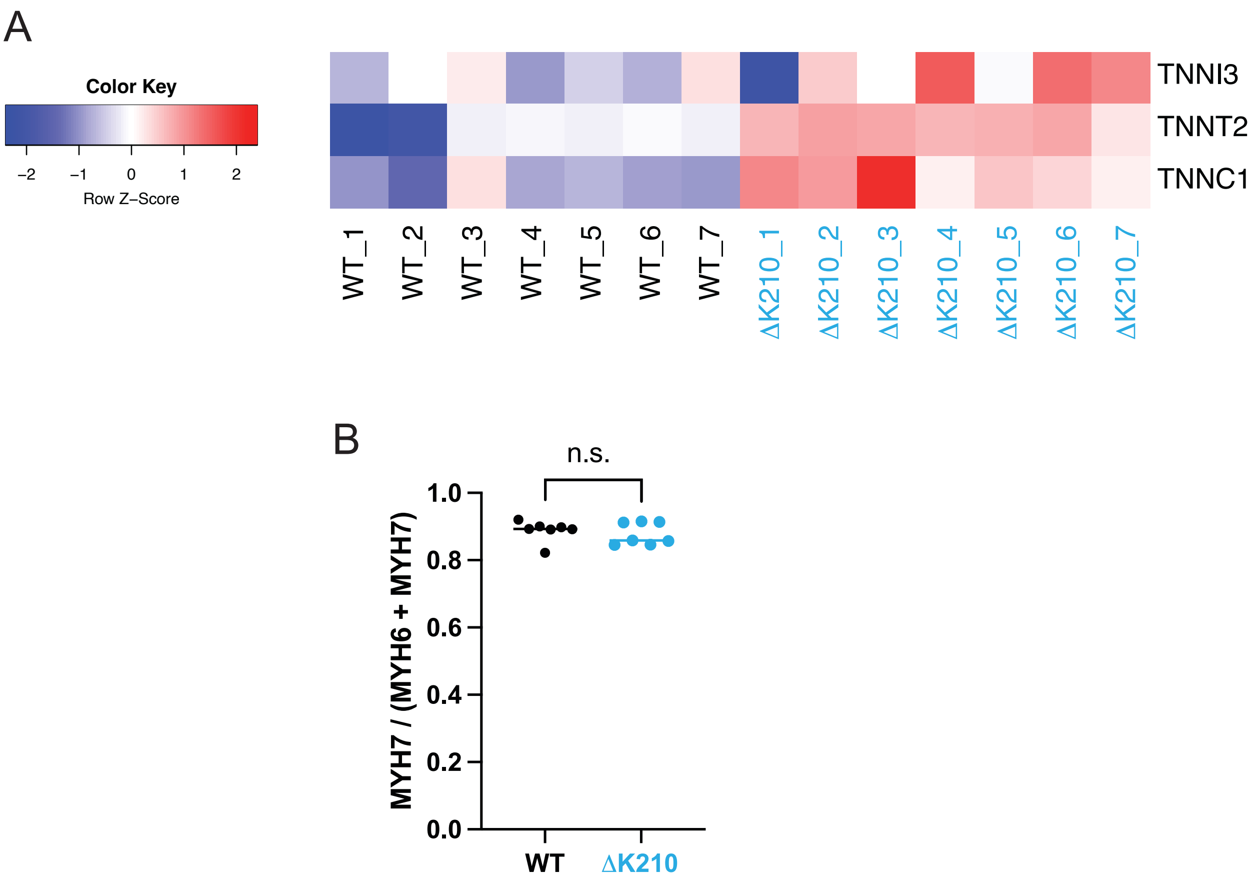


**Supplemental Figure 4: Mass spectrometry measurements of key sarcomeric proteins in engineered heart tissues (EHTs).** (**A**) Expression levels of troponin subunits are increased in hiPSC-CM$^{\Delta K210/WT}$ EHTs relative to WT EHTs. (**B**) MYH7 is the primary myosin heavy chain expressed in EHTs, and the fractional expression of MYH7 is not different between WT and hiPSC-CM$^{\Delta K210/WT}$ EHTs. Bar shows the median and each point represents a separate sample. N = 7 WT and hiPSC-CM$^{\Delta K210/WT}$ EHTs. Statistical testing was done via a 2-tailed T-test.

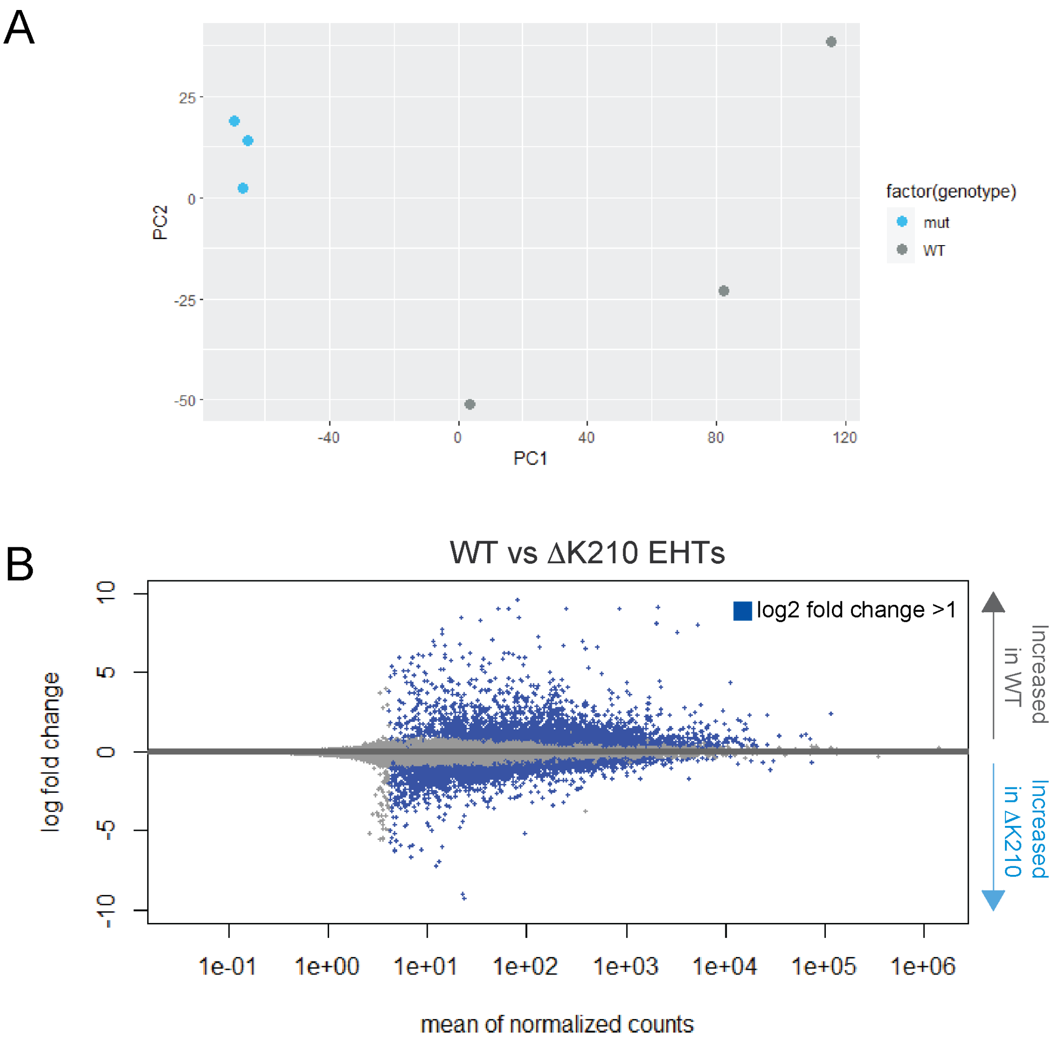


**Supplemental Figure 5: RNAseq data examining differentially expressed genes.** (**A**) PCA plot showing segregation of EHTs with genotype. (**B**) MA plot showing differential expression changes between WT and mutant EHTs.

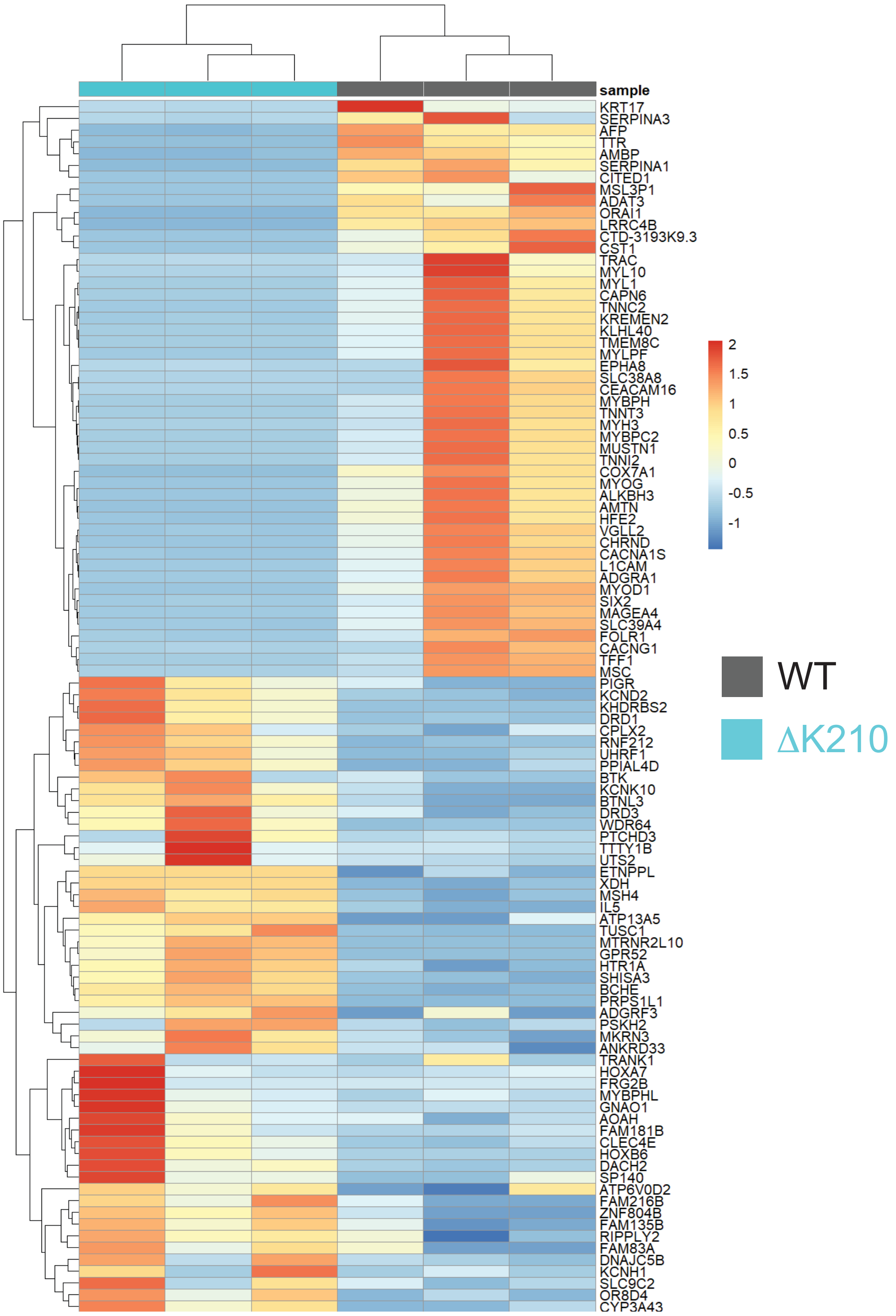


**Supplemental Figure 6: The top 50 genes differentially expressed between WT and hiPSC-CM$^{\Delta K210/WT}$ EHTs.**

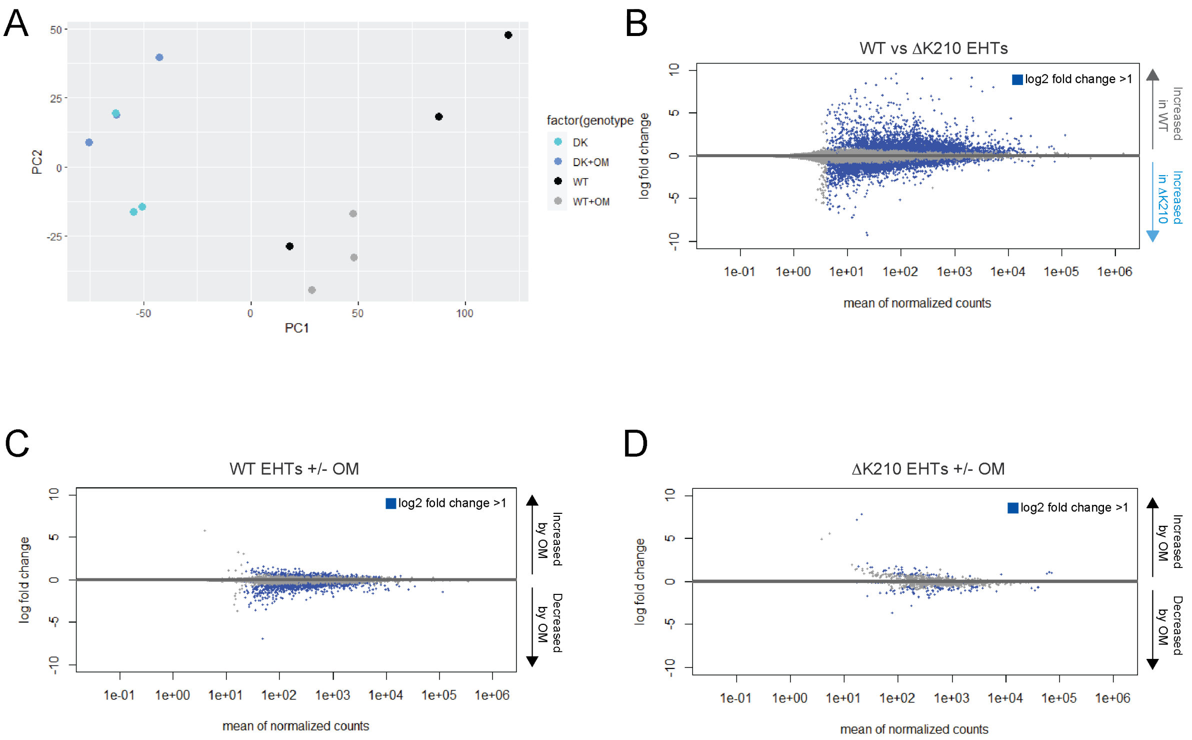


**Supplemental Figure 7: PCA and MA plots for RNAseq of tissues with or without treatment of 250 nM OM for 5 days.** Comparisons are shown for WT, WT + 250 nM OM, ΔK210, and ΔK210 + OM EHTs.

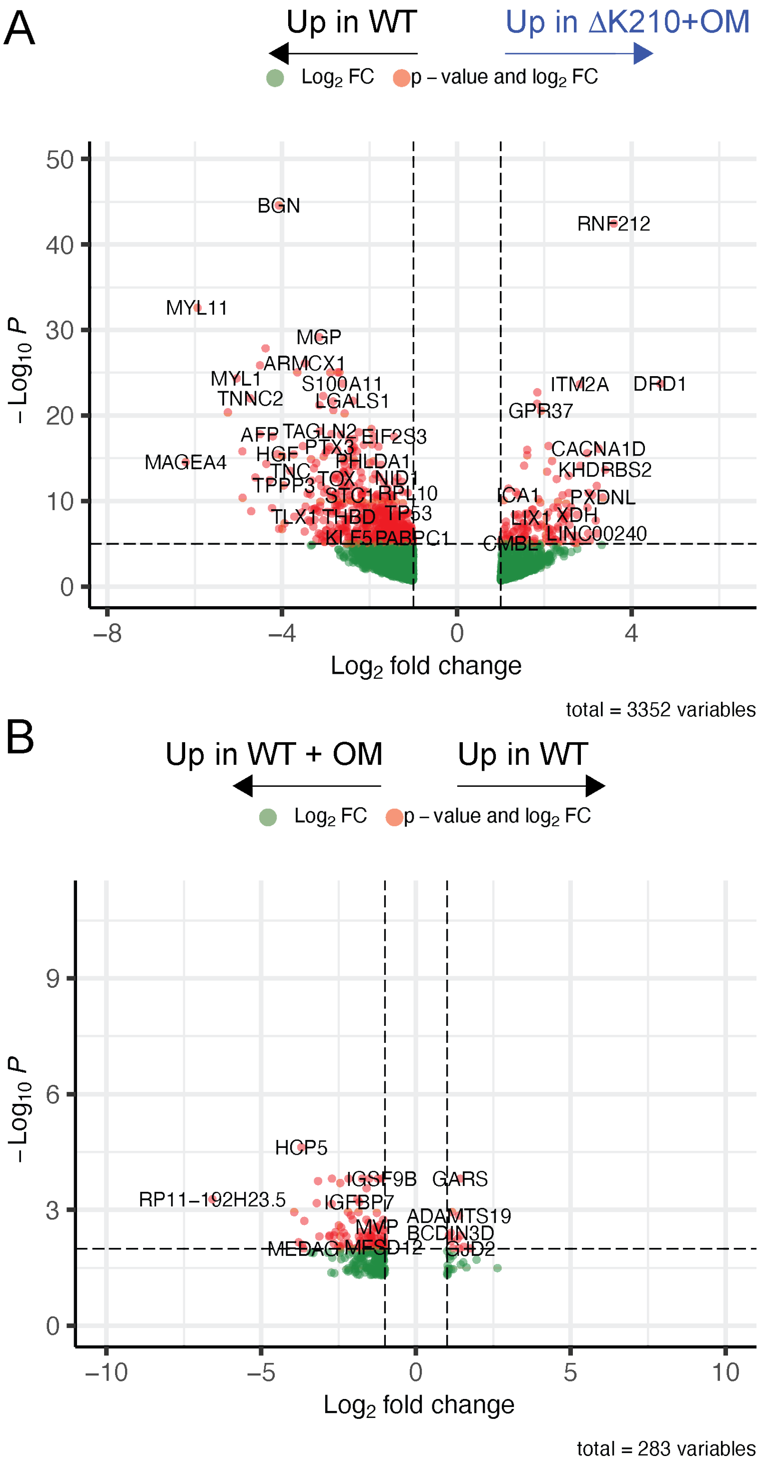


**Supplemental Figure 8: Volcano plots for RNAseq of EHTs treated with 250 nM OM for 5 days.** (**A**) Comparison of ΔK210 + OM and WT – OM EHTs shows that OM does not reverse aberrant gene expression in the mutant. (**B**) Comparison of WT EHTs with and without OM treatment.

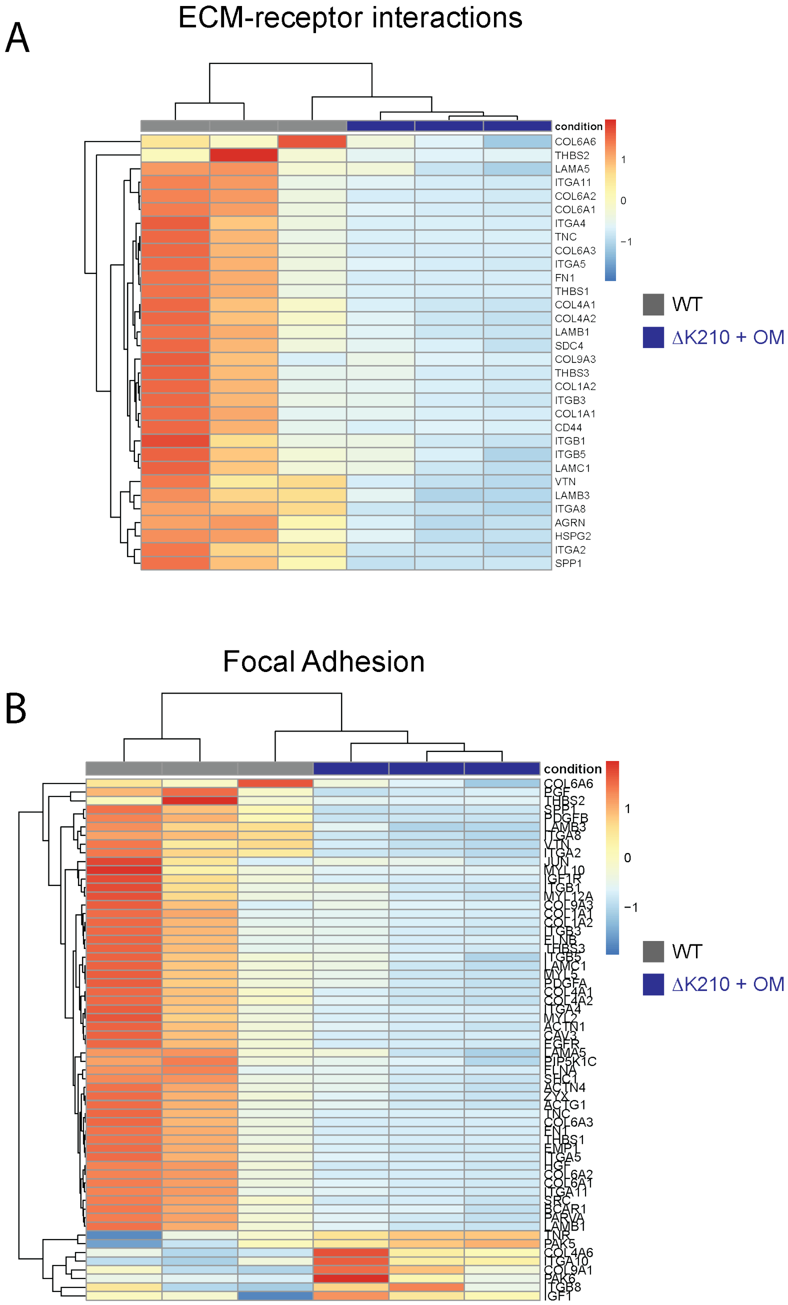


**Supplemental Figure 9: RNA expression of key genes in EHTs demonstrate that 5 days of 250 nM OM treatment does not rescue the expression of key genes in ΔK210.** Dysregulation of genes involved in (**A**) ECM receptor interactions and (**B**) focal adhesions remains after treatment.

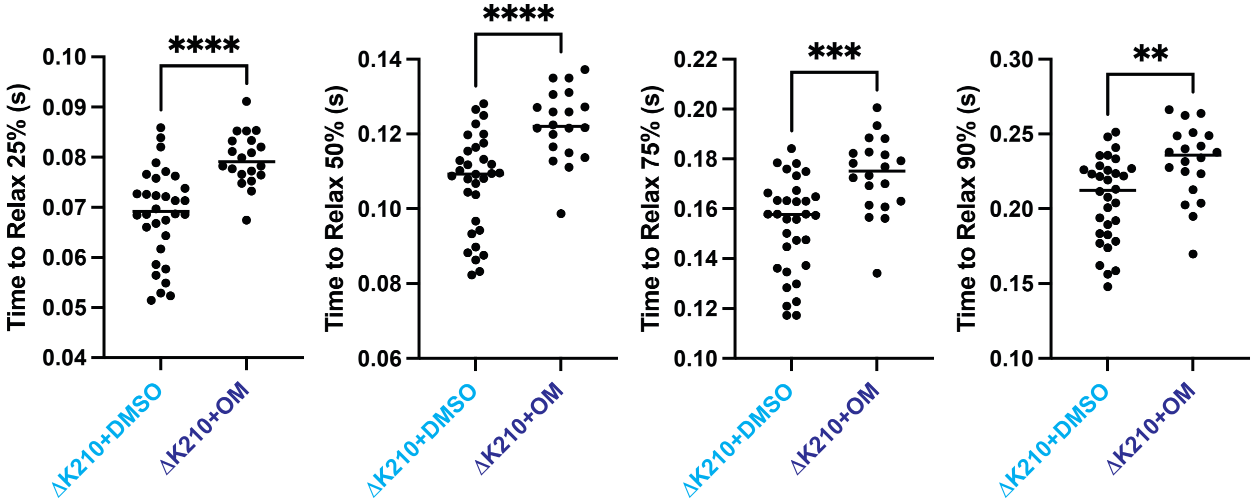


**Supplemental Figure 10: Relaxation times for ring shaped tissue.** N = 32 ΔK210 and N = 20 ΔK210 + OM tissues. Solid bars show median and error bars show interquartile range. Statistical testing was done using a Mann-Whitney test.

### Supplemental Movies

**Supplemental Movie 1: WT EHT formed between two PDMS posts**. Videos were recorded at 100 Hz.

**Supplemental Movie 2: hiPSC-CM$^{\Delta K210/WT}$ EHT formed between two PDMS posts.** Videos were recorded at 100 Hz.

### Supplemental Tables

**Supplemental Table 1: Spreadsheet containing all genes identified by RNAseq.**

**Supplemental Table 2: Spreadsheet containing differential gene expression from RNAseq comparing WT and ΔK210 EHTs.**

**Supplemental Table 3: Spreadsheet containing all proteins identified by mass spectrometry.**

**Supplemental Table 4: Spreadsheets containing differential protein expression from mass spectrometry experiments for WT and ΔK210 EHTs.**

**Supplemental Table 5: Spreadsheet containing differential gene expression from RNAseq comparing WT EHTs with and without OM.**

**Supplemental Table 6: Spreadsheet containing differential gene expression from RNAseq comparing ΔK210 EHTs with and without OM.**
